# Unstable Population Dynamics in Obligate Co-Operators

**DOI:** 10.1101/208934

**Authors:** Abdel Halloway, Margaret A. Malone, Joel S. Brown

## Abstract

Cooperation significantly impacts a species’ population dynamics as individuals choose others to associate with based upon fitness opportunities. Models of these dynamics typically assume that individuals can freely move between groups. Such an assumption works well for facultative co-operators (e.g. flocking birds, schooling fish, and swarming locusts) but less so for obligate co-operators (e.g. canids, cetaceans, and primates). With obligate co-operators, the fitness consequences from associations are stronger compared to facultative co-operators. Consequently, individuals within a group should be more discerning and selective over their associations, rejecting new members and even removing current members. Incorporating such aspects into population models may better reflect obligately cooperative species. In this paper, we create and analyze a model of the population dynamics of obligate co-operators. In our model, a behavioral game determines within-group population dynamics that then spill over into between-group dynamics. Our analysis shows that group number increases when population dynamics are stable, but additional groups lead to unstable population dynamics and an eventual collapse of group numbers. Using a more general analysis, we identify a fundamental mismatch between the stability of the behavioral dynamics and the stability of the population dynamics. When one is stable, the other is not. Our results suggest that group turnover may be inherent to the population dynamics of obligate co-operators. The instability arises from a non-chaotic deterministic process, and such dynamics should be predictable and testable.

## 1. Introduction

Cooperation – where the action of an individual benefits a recipient – is widely observed in nature, from the level of genes to organisms (West et al., 2007; Nowak, 2006). Individuals will choose to cooperate with others if fitness benefits, whether direct or indirect, outweigh the costs (Hamilton, 1964; Trivers, 1971; Taylor, 1992). Because cooperation leads to fitness benefits, it can have significant impacts on a species’ population dynamics as individuals associate with others to form groups for greater fitness benefits. Linking the dynamics at the group level to the entire population will lend a greater understanding of the effects of cooperation on group formation and subsequently overall population dynamics (Bourke, 2011; Bateman et al., 2018).

The type of and context for cooperation underpins group formation and must be considered when understanding the population dynamics of cooperative species. Cooperative societies can be categorized into two broad types: casual and demographic (Wilson, 1975). Casual societies (e.g. starling murmurations or fish schools) are characterized by a constant turnover of group membership with little to no lasting impact on a member’s fitness. Demographic societies (e.g., most social primates, canids, cetaceans, elephants, lions, and eusocial insects), on the other hand, show limited exchange between groups, and dispersal between groups can have lasting fitness impacts on the group’s members. This difference is due to whether cooperation is facultative and therefore helpful but not necessary for survival (casual societies) or obligate where individuals need others for survival and successful reproduction (demographic societies). The difference in cooperation means that obligate cooperators are more selective in their associations and helps explain their limited dispersal. Therefore, one would expect that the differences in cooperation would lead to distinct impacts on their population dynamics, particularly regarding stability.

Typically, swarm dynamics have been used to model the social and population dynamics of cooperative species (Couzin and Krause, 2003; Okubo, 1986; Gueron and Levin, 1995; Zemel and Lubin, 1995; Gueron et al., 1996; Bonabeau et al., 1999; Mirabet et al., 2007; Saffre and Deneubourg, 2002). This approach has worked well to model the population dynamics of facultatively cooperative species, however not as well for obligately cooperative species. While actors in a swarm model are selective regarding associations, dispersal is generally not limited with individuals moving freely between groups. On the other hand, some models with limited dispersal do not permit selectivity of any individual (Courchamp et al., 1999; Courchamp et al., 2000; Dennis, 2002). And more generally, not all individuals are selective in population models of cooperators. Whether models of swarm dynamics or of limited dispersal, the focus remains on an individual disperser and their choice of associations with non-dispersers lacking choice (Lehmann et al., 2006; Parvinen and Brannstrom, 2016). Including both limited dispersal and group-wide selectivity into a model of cooperative species’ dynamics should better reflect the dynamics of obligate co-operators.

In this paper, we model the population dynamics of obligately cooperative species. First, we construct and analyze a specific model of population dynamics embedded with a behavioral game with group-wide selectivity. In this model, there is within-group cooperation, within-group competition, between-group competition, and limited dispersal between groups (specifically no movement). We analyze the model’s population dynamics and behavioral dynamics separately before combining the two. From there, we relax our assumptions and generalize our analysis to a class of models that keeps all key features of the specific model. We show a mismatch in stability: a stable population size is not behaviorally stable and groups that are behaviorally stable are unstable in terms of population dynamics. In other words, a group can either have behavioral stability or population size stability but never both. We go on to discuss the implications of our model including its relevance to real systems and future applications.

## 2. Dynamics of the Specific Model

### 2.1. Population Dynamics

To model the population dynamics of obligate cooperators, we first assume that all individuals are cooperative and do so by forming associations, i.e. groups, to gain greater fitness. Here, we define fitness as an individual’s expected per-capita growth rate. For the sake of simplicity, we assume all individuals are identical, distinguished only by whether they are within or outside a group which gives all individuals within a group the same fitness function. While individuals are discrete, for analysis we treat the set of individuals and the corresponding fitness function as continuous. While this is mostly for mathematical tractability, representing individuals as a continuous set works when the number of individuals is extremely large or represents density. Furthermore, the continuous form of the equation can turn discrete through rounding.

When individuals associate, an emergent group-level cooperation appears as the aggregation of each individual’s cooperative acts (Dugatkin, 1998). In terms of population dynamics, this is often modeled as an Allee effect, or the positive relationship between an individual’s fitness and its associations within a group – typically measured as group size (Allee, 1931; Allee, 1938; Trivers, 1971; Axelrod and Hamilton, 1981; Dugatkin, 1998; Nowak, 2006; Stephens et al., 1999; Angulo et al., 2018). We include this Allee effect to within-group dynamics but not between-group dynamics. We imagine that individuals benefit from group living via collective foraging, defense, or other positive social interactions yet suffer from resource sharing, disease transmission and other negative interactions. Thus, groups are akin to habitat sites with an Allee effect. Specifically with group size, we assume that benefits to an individual increase linearly while fitness costs increase super-linearly leading to an overall humped-shaped relationship between group size and fitness (Terborgh, 1983). We believe this relationship between group size and fitness is biologically relevant because there are limits to growth within a group. We considered alternate forms, such as increasing fitness with group size with diminishing returns. However, in the absence of competition, these would result in infinitely sized groups. Regardless, all forms should have a broadly similar overall result on stability (see Discussion, The Limits to the Model).

Our model focuses on the process of group creation and formation of obligate cooperators (Bourke, 2011). In facultative cooperation, individuals move more freely between groups which means that migration between groups is a bigger determinant of a focal group’s size than fitness of individuals within that group. As such, models of group formation in which fitness at the individual level immediately translates to overall population size while group size is the division of the overall population work well for facultative cooperators. However, these models work less well when dispersal and migration is limited. This is because the fitness of individuals within a group is a bigger determinant to that group’s size than migration. As such, when modelling the population dynamics of obligate cooperators, we model the fitness dynamics of each individual group then aggregate those dynamics to create the overall population dynamic.

With our assumptions, we imagine *n*_*g*_ different groups, each with their own group size. The fitness function (per capita growth rate) for a focal group *i* is given by equation (1).

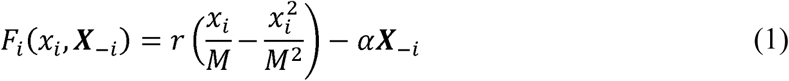

Here, *x*_*i*_ is the size of focal group *i*, ***X***_*−i*_ is the cumulative size of all other groups *∑*_*j≠1*_ *x*_*j*_, *r* is a growth rate scaling factor, *M* is maximum potential group size, and α is the strength of intergroup competition (potentially determined by the depletion of shared resources). Since we assume all individuals are identical, all groups have the identical parameters allowing us to rescale per-capita growth rates by *r* to give:

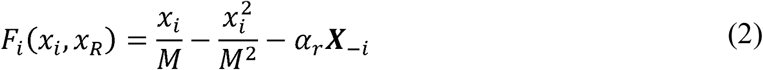

where 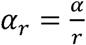 is the ratio of intergroup competition to the growth rate scaling factor. Assuming that is held constant, we can denote the equilibrium group size of a single group as 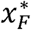 where growth rate equals zero. Solving for 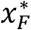, we obtain two values where 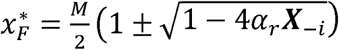.

The first two terms of equation (2) define these intra-group dynamics where the first term represents the fitness benefits of group living and the second term represents the fitness costs of group living. The third term of equation (2) represents the loss of fitness due to inter-group competition and is broadly assumed to be linear. In equation 2, fitness at first increases at low group sizes 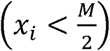 and then decreases at high group sizes 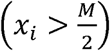, and such within-group interactions create an Allee effect (Fig. 1b) (Allee, 1931; Allee, 1938). The strength of any Allee effect is typically classified as strong or weak based upon whether or not there is an extinction threshold respectively (Courchamp et al., 1999b; Stephens et al., 1999; Angulo et al., 2018). In our model, the strength of the Allee effect is flexible and depends on the presence or absence of competition external to the group. With intergroup competition, the Allee effect is strong. Conversely, without intergroup competition, the Allee effect is weak.

**Fig. 1.**
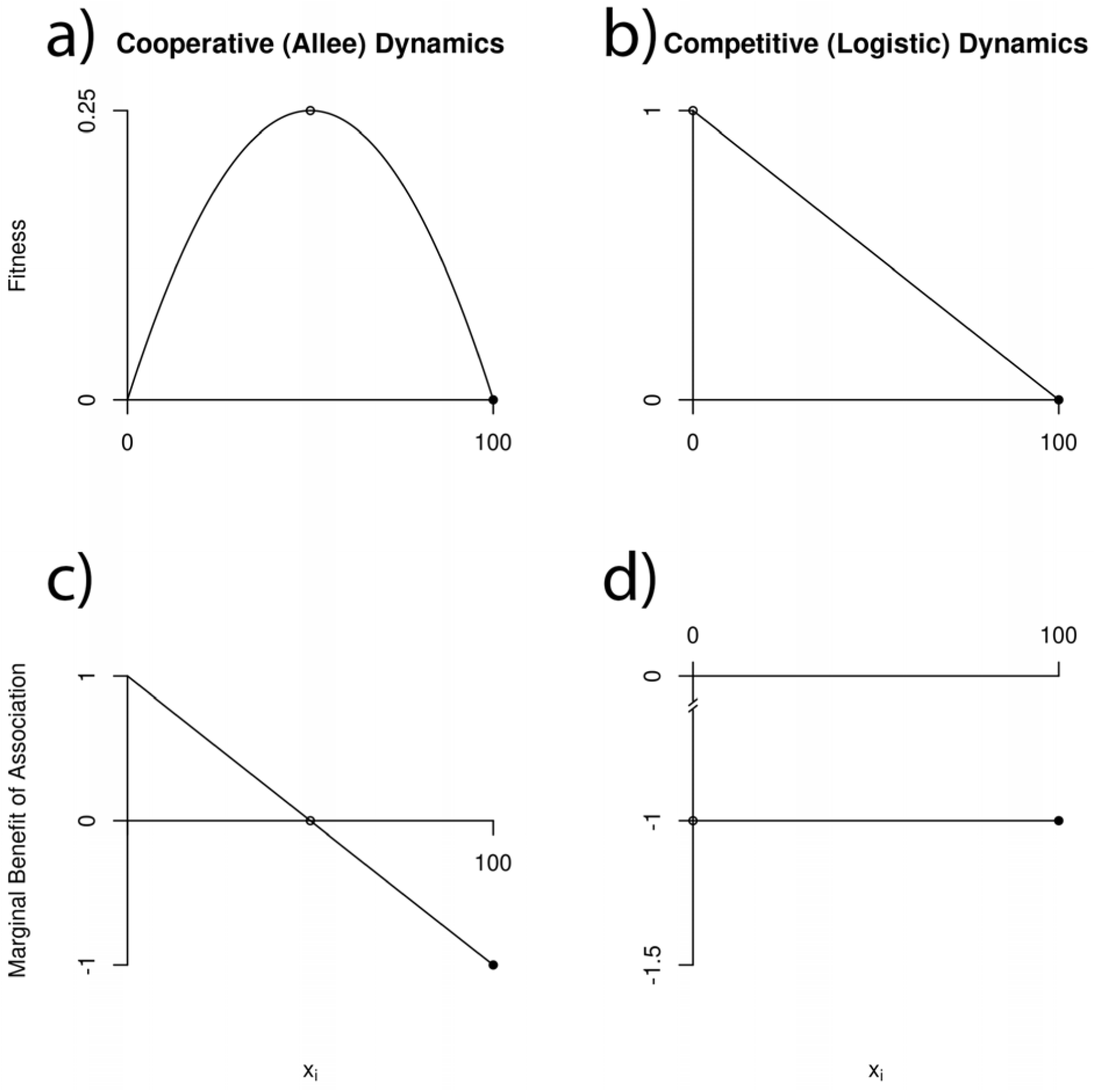
A figure of the fitness (a, b) and association (c, d) functions of our model (a, c) and strictly competitive model 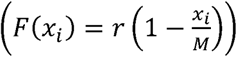 (b, d). (a) Our model’s fitness function (b) The competitive model’s fitness function (c) Our model’s association function (d) The competitive model’s association function. Open circles indicate the group size that gives the maximum fitness, i.e. optimal group size, while closed circles indicate the stable fitness equilibrium. One can see that in both systems, the optimal group size and stable fitness equilibrium do not match; however, there is greater implication on the population dynamics in our model as the association function has both positive and negative elements, lending itself to behavioral game. *r*= 1,*M* = 100.

We can now analyze the overall population dynamics of the entire system of groups. This analysis replicates Wang et al. (1999), revealing the same qualitative results with minor differences. Therefore, we keep this section brief. With a single group, there are two equilibria: 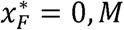. The first equilibrium is unstable, the second is stable. Therefore, any group of strictly positive size will grow or shrink to *M* (Fig. 2a,b,c). The addition of another group changes the dynamics since it introduces a positive ***X***_*−i*_ which then varies based on the size of the other group. Outcomes now depend upon the strength of inter-group competition *α_r_* (Fig. 3). If inter-group competition is strong 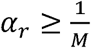, then either only one group survives or both groups go extinct (Fig. 3a). The equilibrium where both go extinct is a saddle-point (unstable except for when the initial population sizes of the two groups are equal) while the equilibria where only one group survives are stable (the group with the larger size outcompetes the other). As inter-group competition weakens 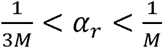, there arises an unstable (saddle point) interior equilibrium (Fig. 3b,c). The equilibrium where both groups go extinct is now fully unstable. The equilibria with just one surviving group remain locally stable.

**Fig. 2.**
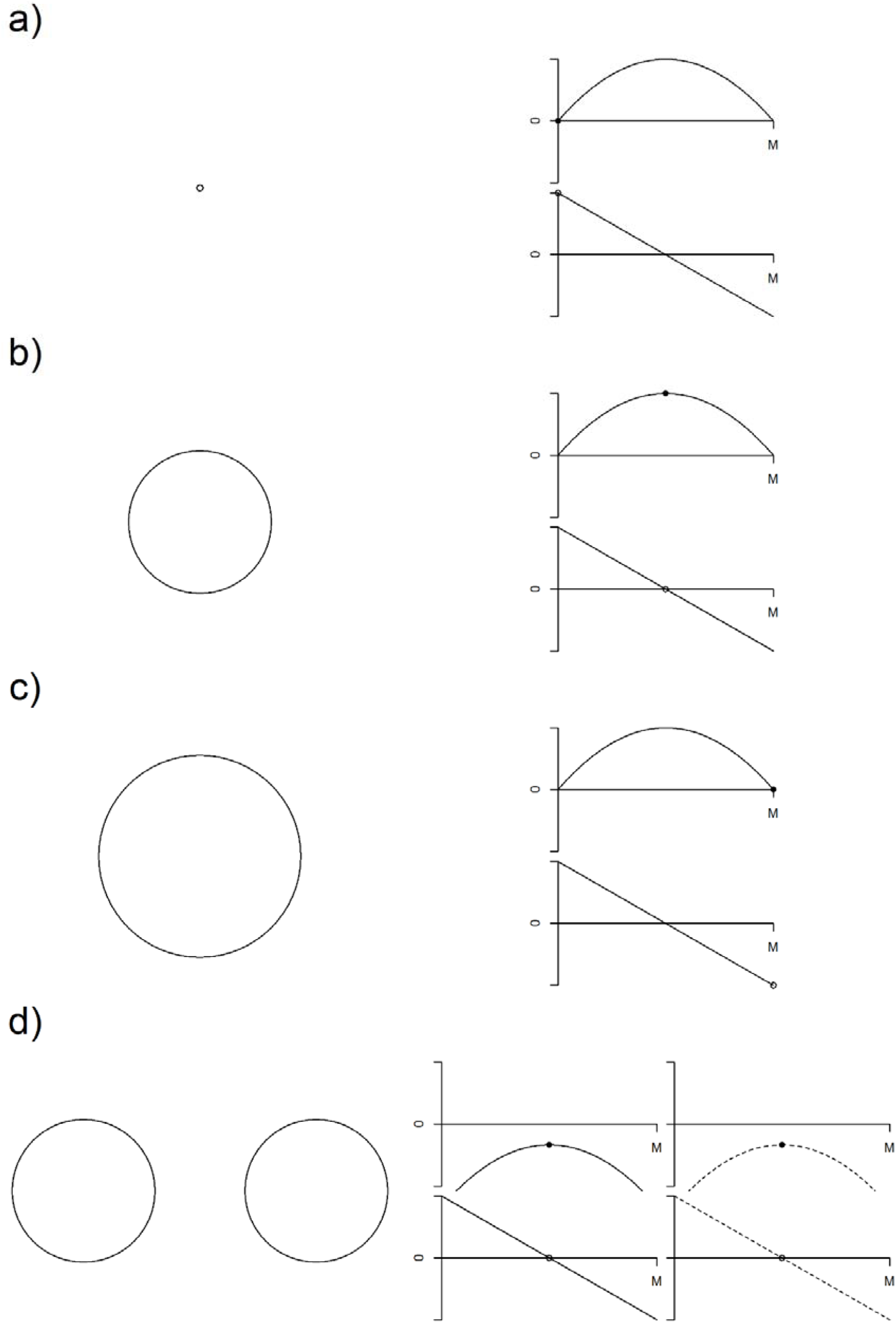
A schematic of the process of group growth and splitting. On the left, a circle represents a group. On the right is said group’s fitness function on top and association on the bottom. Our parameters for this model are *r* = 1,*M* = 100, 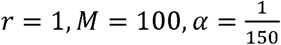. (a) The fitness function of a single group at extremely small population size *x*_1_ = 0.5. (b) The fitness function of the group when it is at its optimal size 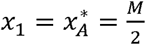. In this case, all members of the group are at maximum fitness and satisfied with group size, but fitness is positive causing the group to continue growing. (c) At this point, the group is at maximum size *x*_1_=*M* so fitness is 0 and it will stop growing, but A(*x*_1_)<0 so the group members are unhappy. (d) The group just after splitting into two groups 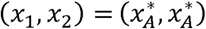. The new group (dotted line) leads to a lowering of fitness due to intergroup competition, which in this case is below 0. Here, *T*_1_ < α_*r*_ < *T*_2_, so both groups will go to an unstable equilibrium.

**Fig. 3.**
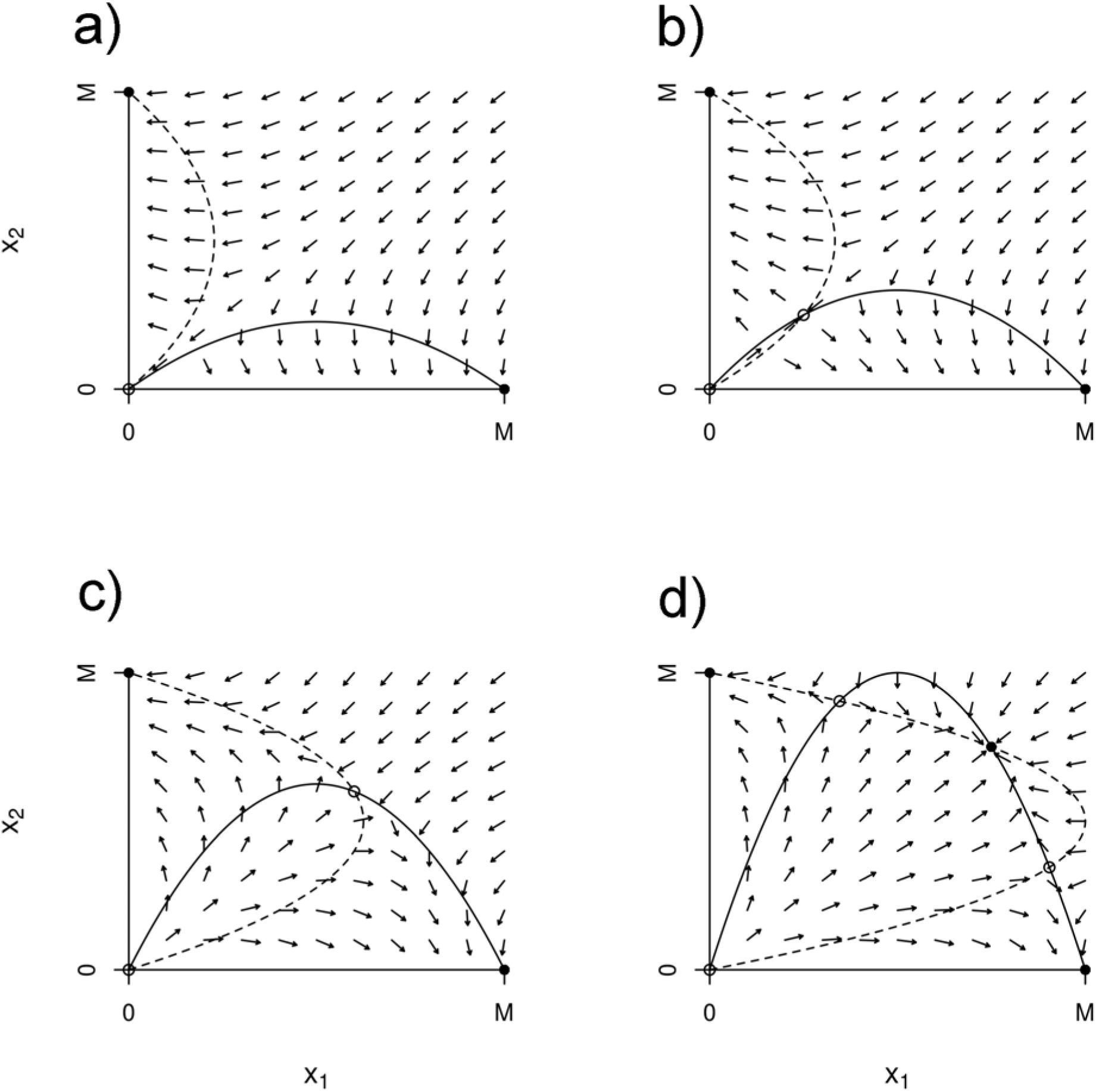
The isoclines, equilibria, and directional field of a two-group system under different strength of competition α_*r*_. (a) α_*r*_ = 1.1/*M* (b) α_*r*_ = 0.75/*M* (c) α_*r*_ = 0.4/*M* (d) α_*r*_ = 0.25/*M*. Solid dots represent stable equilibria while open dots represent unstable equilibria.

If competition weakens even further 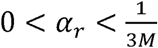, two new interior equilibria appear where both groups co-exist at positive but unequal sizes. These two new interior equilibria are unstable. The former interior equilibrium where both groups have the same population size now becomes locally stable (Fig. 3d). With weak competition, there is a strong Allee effect and each group has an extinction threshold based on the number of individuals in the other group. If the initial sizes of both groups are above their extinction thresholds, competition is too weak to drive extinction, and the two groups will persist at the interior equilibrium of equal sizes. If one or both groups are below their extinction thresholds, then the smaller group be extirpated and the larger group will grow to size *M*. As intergroup competition disappears, the interior equilibrium with equal sizes becomes globally stable as both reach maximal carrying capacity, and the other interior equilibria merge with the two in which only one group survives. These two equilibria and the one in which both groups go extinct are all unstable.

As we add additional groups, the same fundamental dynamics remain (see SI). The only difference is that inter-group competition must be weaker – specifically, it must be scaled by 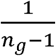 where *n*_*g*_ is the number of groups – for the interior equilibrium to be locally stable. More generally, we can say that for any system of *n*_*g*_ groups, the only stable interior equilibrium (all groups at strictly positive size) occurs when all groups, equal in size, are at a size 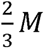 or greater. For this to occur with an increasing number of groups, inter-group competition must decrease. If not, then the addition of more groups destabilizes the interior equilibrium and leads to a collapse of group numbers.

### 2.2. Behavioral Dynamics

Embedded in equation 2 is a behavioral dynamic. Changes in group size lead to changes in fitness as defined as per-capita growth rate. If we take the partial derivative of the fitness function with to population size, we obtain a function (hereafter called the association function) *A*_*i*_(*x*_*i*_) that indicates the marginal contribution of an individual to the fitness of other members in group *i* (equation 3).

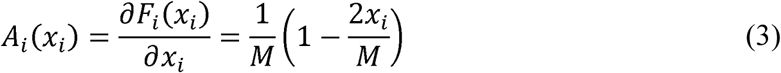

Assuming that individuals are rational and seek to maximize their fitness, they should want to be in a group whose size maximizes their fitness. If individuals have limited information – specifically that they can only “see” their fitness in relation to the size of their current group and are blind to the influences of non-group members – then this association function guides their choices regarding optimal group size. We call the group size which maximizes fitness the optimal group size and denote it by 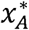. For this equation, optimal group size is half of maximal group size 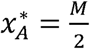.

Typically in models of cooperative species, only the dispersing individual is selective and given choice of association. However, all individuals in a group should be just as selective as the disperser. Because we do not include a specific mechanism of cooperation and simply assume that all individuals cooperate, we can use coalitional (a.k.a. cooperative) game theory to analyze the behavior of the individuals within a group en masse. Coalition game theory seeks to understand two things: how individuals select their associations and how the resulting payoff to that coalition is distributed in an equitable and efficient manner (von Neumann and Morgenstern, 1944; Peleg and Sudhölter, 2007). In coalitional game theory, there are *N* players who can choose to associate with other players to form various coalitions 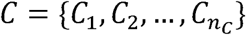. Through their associations, they receive a payoff *v*(*C*_*i*_). If all players choose to form a single coalition *C* = {*N*}, this is known as the grand coalition. In our case, the players are the individuals within a group *N*=*x*_*i*_, and the total payoff to the group is its growth rate *v*(*C*_*i*_)= *x*_*i*_ *F*_*i*_(*x*_*i*_). By assuming that all individuals are identical, payoffs are equally shared among all members and can be represented as fitness or per-capita growth rate *F*_*i*_(*x*_*i*_). Using this framework, we can analyze how individuals should choose their associations.

For illustrative purposes, we analyze the behavioral game in which there is a single group facing no inter-group competition (α_*r*_ ***X***_*−i*_ = 0) However, the general principles remain the same regardless of the presence or strength of inter-group competition. Based on the association function *A*_*i*_(*x*_*i*_), the marginal contribution of an individual is positive *A*_*i*_(*x*_*i*_) > 0 when group size is less than the optimal size 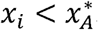. Under these conditions, the game is super-additive, i.e. all individuals receive greater fitness from being in one large coalition of *x*_*i*_ rather than separated into smaller coalitions. As such, there is no incentive for any individual to seek a group of a lower size. However, if the group is larger than its optimal size 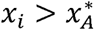, then the behavioral game is no longer super additive. Instead, the marginal cost of an individual is negative *A*_*i*_(*x*_*i*_)< 0. Therefore, individuals in a marginally smaller group will obtain greater fitness. As such, individuals in a group of size 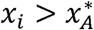 will prefer to be in a smaller coalition, specifically at the optimal group size 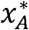 (Fig. 2b). The group is not behaviorally stable as individuals will leave or be forced out from the current group, creating new groups in the process (Fig. 2). Summarily, through coalition game theory any group smaller than optimal group size is behaviorally stable while any group larger than optimal group size is behaviorally unstable.

Now with population and behavioral dynamics analyzed separately, we can combine them to derive a full picture of the population dynamics of cooperative species. We seek the possibility of a joint population and behavioral equilibrium and show it to be impossible.

### 2.3. Overall Population Dynamics

In our specific model, any fully stable equilibrium must have all groups at a size 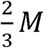 or greater and that more groups in a system lead to greater instability. Analysis of behavioral dynamics shows that current groups will split and new groups will be formed so long as the current group’s size is beyond 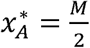. Combining the two results, we obtain the main result: over time, the behavioral dynamics of cooperative species will tend towards more groups when population dynamics are stable, and a stable behavioral equilibrium will lead to unstable population dynamics and an eventual collapse of group numbers. Through this, we see that there is a mismatch between the regions where population dynamics and behavioral dynamics are stable. Later, we show this to be a more general phenomenon. For now, we look in detail at the potential dynamics of our specific model.

We add to our model the limited dispersal behavior seen in obligate cooperators. To do so, we make additional behavioral assumptions: groups can only split (i.e. members can leave or be forced out of a group but not join other groups) and group size only increases through reproduction (a consequence of the former assumption). In addition, we assume that behavioral dynamics only occur when all groups have reached a fitness equilibrium, i.e. group splitting only occurs at the fitness equilibrium. We recognize that more efficient behavioral dynamics could occur with members being forced out once an optimal group size has been reached. We chose our assumption as it gives the resulting model analytical tractability. Furthermore, some cooperative species split and show mass dispersal behavior at group sizes beyond the behavioral optimum, e.g. bees, rhesus macaques, and orcas (Fell et al., 1977; Dittus, 1998; Stredulinsky, 2016).

In our analysis, we begin by assuming a single group without competition. So long as initial population size is strictly positive, the group will either grow or shrink to the stable equilibrium 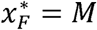. Our behavioral analysis shows that a single group of size *M* is unstable. There is a strong incentive for the members to achieve a group of size 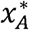. Therefore, as the group’s size reaches 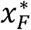, it splits into two groups, both of size 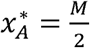 (Fig. 2d).

After the split, the addition of another group means both groups are facing competition with ***X***_*−i*_ equaling 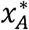 for both groups. The fitness for both groups is 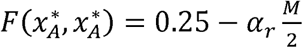. The size of inter-group competition *α*_*r*_ will determine the long-term dynamics of our system. There are two thresholds of *α*_*r*_. There is the main threshold *α*_*r*_ = *T*_1_ which divides competition into strong and weak, and a secondary threshold *α*_*r*_ = *T*_2_ > *T*_1_ which divides strong competition further into moderately strong and extremely strong. These three strengths of competition – extremely strong, moderately strong, and weak – correspond to the dynamics of total extinction, unstable equilibria, and group turnover respectively (Fig. 4).

**Fig. 4.**
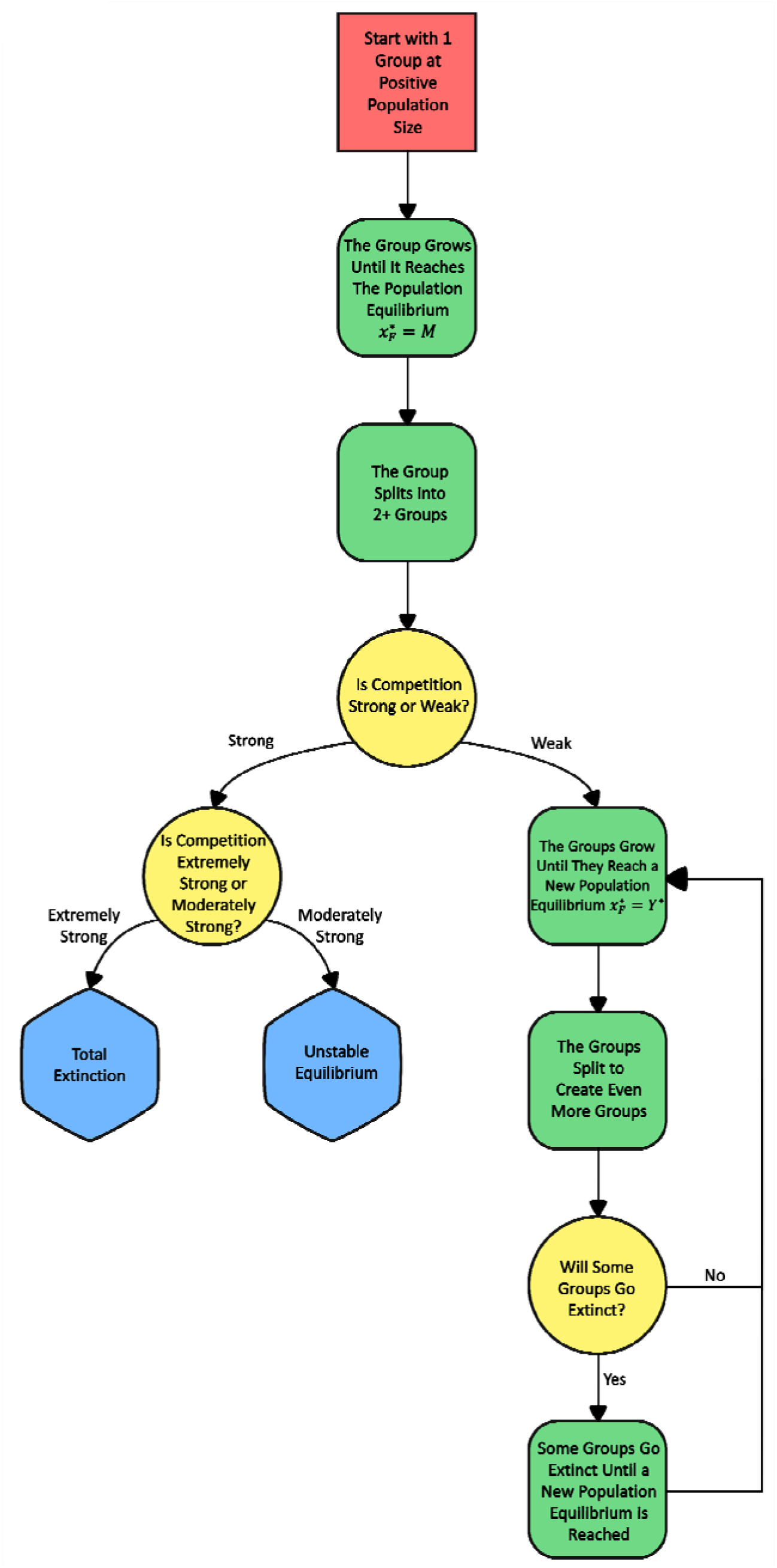
A flowchart for how our model evolves starting with a single, cooperative group. Red-filled squares represent initial conditions, green-filled rounded rectangles represent transition states, yellow-filled circles represent “decision” points, and blue-filled hexagons represent end points. As seen here, there are either no endpoints (weak competition) or the endpoints are unstable (strong competition).

If competition is extremely strong, *α*_*r*_ > *T*_2_ > *T*_1_, then competition between the two groups is so severe as to drive both groups to extinction, i.e. there is no positive population size at which both groups can persist. If competition is moderately strong *T*_1_ < *α*_*r*_ < *T*_2_, then competition between the two groups is strong enough to reduce population size but not strong enough to drive the group to extinction. Therefore, both groups come to rest at a positive population size that is smaller than 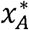. As previously shown in the section on fitness dynamics, the equilibrium is unstable with respect to population (Fig. 3b). These results, while biologically different, are mathematically similar. When competition is weak, *α*_*r*_ < *T*_1_ < *T*_2_, then an entirely different dynamic occurs. Since fitness is positive when both groups are at 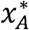, both groups will continue to grow to some new fitness equilibrium 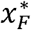. Because the new fitness equilibrium is larger than optimal group size 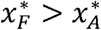, the two groups will split to become four, two larger at 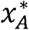 and two smaller. The two smaller groups will either shrink and be extirpated and reset the system to the previous state or persist with the result being four groups at a new fitness equilibrium 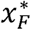. At this point, the same process of the two groups will happen with the four groups. Whether at two, four, or some other number of groups, there will be a population size such that new groups will be extirpated returning the system to the previous state and leading to a turnover of new groups (Fig. 5).

**Fig. 5.**
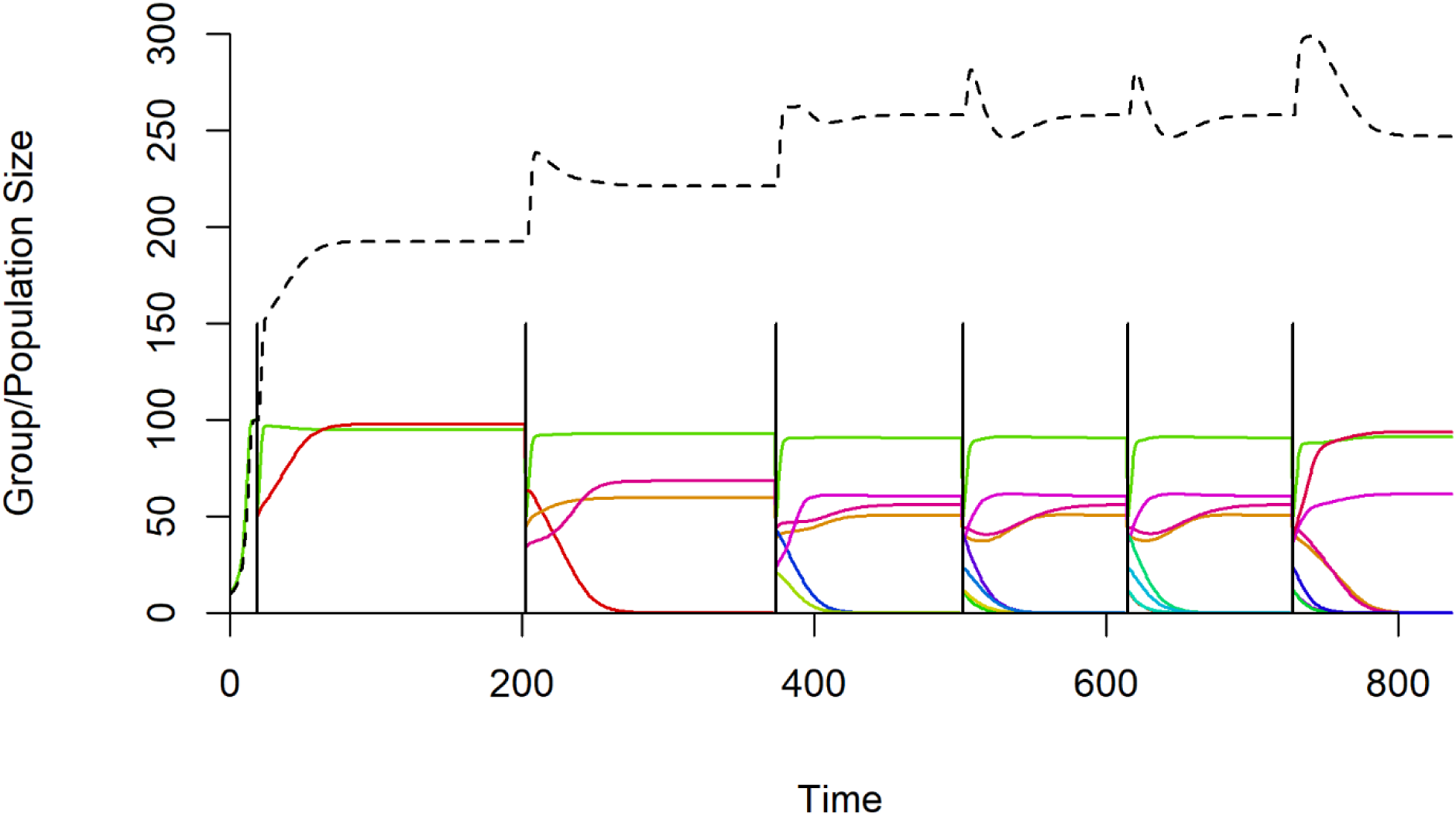
A time series of the populations when 0< *α*_*r*_ < *T*_1_. One can see an initial buildup of groups and overall population before transitioning to group turnover and “oscillations”. Each group is represented by a color with groups constantly appearing, shrinking, and going extinct. Each new group was given a new *r* and *M* based on the logit normal distribution. The large, solid, vertical, black lines represent a time when the existing groups split. The dashed line represents the total population size.

Further derivation of these three dynamics can be found in the supplementary information.

## 3. Argument for General Instability of the Behavioral and Population Dynamics

Now we present a more general demonstration of the fundamental mismatch between a stable population equilibrium and a stable behavioral equilibrium. We retain the majority of our earlier fundamental assumptions including 1) cooperation leading to fitness benefits, 2) all individuals being identical and only distinguished by presence or absence in a group, 3) individuals seeking to maximize their fitness but only know the relationship between the size of their current group and their fitness. However, we relax our previous assumptions relating to dispersal. Individuals can join existing groups, and behavioral dynamics can occur concurrently with the population dynamics.

Let 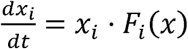 be the population dynamics within group *i* where *F*_*i*_(*x*) gives the expected per capita growth rate and is assumed to be a continuously differentiable (at least *C*^2^) function of 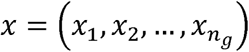 which is the vector of group sizes for all groups numbering 1 to *n*_*g*_. We refer to *F*_*i*_(*x*) as group *i*’s fitness function. Taking the partial derivative of 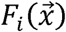 with respect to *x*_*i*_ and fixing all other variables to the 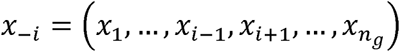 results in a function of association 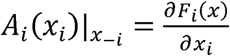 for focal group *i* which gives the change of the group’s per capita growth rate due to additional members, i.e. an individual’s marginal fitness contribution. When 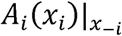 is positive (or negative), additional members increase (or decrease) an individual’s fitness. We call these states cooperative and competitive respectively. 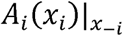 also determines the preference of individual group members for more or fewer members. In a cooperative state, individuals at best do not want the group size to decrease and will resist splitting and behaviorally favor an increase, and vice-versa for group sizes in a competitive state.

Let 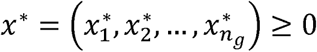 be a solution to the equation *F*_*i*_(*x\**) = 0 for all *i* ∈ {1, … *n*_*g*_}.Let *i*^+^ ⊆ {1, … *n*_*g*_} be the subset of all groups with positive population size. We can analyze the stability of this point through the Jacobian *J*. The diagonals of the Jacobian are 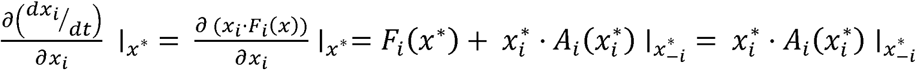 for all *i* ∈ {1,2, … *n*_*g*_}. As 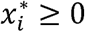, the diagonals are either 0 or reflect the sign of the behavioral game at that point. If the equilibrium is cooperative for all groups in the set 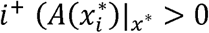 for all *i* ∈ *i*^+^, then the trace of the Jacobian is positive, Tr(*J*) > 0. Since the sum of all eigenvalues is positive, at least one eigenvalue is positive, and this vector of equilibrium population sizes is unstable.

If at least one of the groups in *i*+ is at a competitive state, then all eigenvalues could be negative, meaning the population dynamics could be at a stable equilibrium. Those competitive groups though are at a behaviorally unstable equilibrium. Since *F*_*i*_(*x*) is *C*^2^ smooth for all *i*,, then there exists a point of lower population size 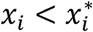 that gives higher fitness 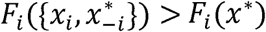. If this is the case, coalition game theory tells us that a coalition of 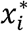 in group *i* will not form; instead, individuals will break off to form a group of size 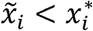 where 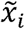 is a group size that maximizes the group’s per capita growth rate.

An individual’s preferred state is one that maximizes its fitness. A necessary condition is that 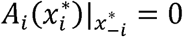 and 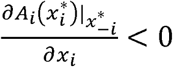. If there is an *x*\* satisfying these conditions for all groups, then diagonals of the Jacobian matrix *J* are all 0; therefore, the sum of all eigenvalues are 0. If this is the case, then there is either a mix of positive and negative eigenvalues (meaning unstable population dynamics) or all eigenvalues are 0. Because 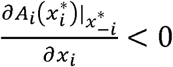, the latter corresponds to a partially stable state and not a neutrally stable state. This means there are clear domains of instability on whose boundary the point *x*\* resides.

According to this analysis, there will be a fundamental mismatch between stable behavioral dynamics and stable population dynamics in obligate cooperative species, and their overall population dynamics will always be unstable.

## 4. Discussion

We have analyzed the population dynamics of obligately cooperative species with limited dispersal. We embedded a behavioral game with these properties into a model of population dynamics to understand group formation and interaction (Bourke, 2011). There were three potential outcomes: 1) extinction of all groups, 2) unstable equilibria of population sizes within groups, and 3) group turnover. The cooperative species fails to achieve stable population dynamics due to a mismatch between the stability of the behavioral equilibrium and that of the population dynamics. While we are not the first to note that local extinctions and extirpations occur in population dynamics due to Allee effects, our model shows them to be intrinsic and unavoidable. Our findings join other mathematical analyses in generalizing the phenomenon to factors intrinsic to obligately cooperative species.

Empirically, many obligately cooperative species including mole-rats, social spiders, and banded mongooses do not show stable population dynamics and instead show constant group turnover (Jarvis et al., 1994; Aviles, 1997; Clutton-Brock et al., 1999). Hypotheses with mathematical support have been developed to explain the phenomenon. Aviles (1999) noted that cooperation can magnify reproductive output, leading to oscillations and chaotic behavior, and ultimately extinction of a group. Chourchamp et al. (1999a) and Wang et al. (1999) both showed the importance of a within-group extinction threshold. Our results further indicate the importance of extinction thresholds. However, whereas groups by assumption had extinction thresholds in prior models, we show that this extinction threshold necessarily emerges due to intergroup competition regardless of whether the Allee effect is strong or weak. Furthermore, more groups lead to greater instability. The models of Chourchamp et al. (1999a) and Wang et al. (1999) relied on external factors to drive the instability. With the addition of a behavioral game of group splitting and formation, we show that new groups will constantly be created, leading to unstable population dynamics and an eventual collapse of group numbers. Among all the hypotheses for the instability of obligately cooperative systems, our model with an embedded behavioral game shows how these dynamics are not only intrinsic but unavoidable.

Additionally, our results show that constant group turnover arises from non-chaotic deterministic interactions. This means that the localized group extinctions are a general, repeatable, and predictable pattern against which field studies and data can be tested. Using simulations and controlled experiments, we can now predict how attributes and traits of species along with environmental variables can affect the cooperative species’ population dynamics (see Future Directions).

### 4.1. Short-Term Intragroup Dynamics

In our model, each group has its own population dynamic with an Allee effect. With the Allee effect, there is a non-zero optimal group size which maximizes fitness for the group members. We assume that individuals in a group whose size is beyond the behavioral optimum will split off to form their own smaller groups. This enhances overall fitness of the group. We are not the first to understand that group splitting can occur due to an Allee effect. For example, Crema (2014) incorporated the Allee effect into a simulation model to understand human settlement dynamics. Fission-fusion group dynamics, permanent or otherwise, are a well-studied aspect of cooperative societies with examples ranging from ants to cetaceans to humans.

In our model, we made some simplifying assumptions at the expense of realism. For both models, we assumed: 1) cooperation leads to fitness benefits, 2) individuals are identical and only distinguished by their present group size, 3) individuals seek to maximize fitness, 4) the overall number of players within a group changes according to the average fitness of group members, and 5) individuals have limited information, specifically they only know about the fitness function of their present group. For the specific model, we added the assumptions that 6) individuals cannot leave a group and join another established group, i.e. groups form by the fission of existing ones, and 7) behavioral dynamics occur when the system has reached the population equilibrium. Some may have more relevance (1, 3, 4) to biological systems than others (2, 6, 7), but these assumptions allow for conclusions to be derived analytically. In most eusocial insects for example, individuals within a caste are largely identical (though not between castes), and in honey bees, creation of a new colony happens when the hive has reached its maximum capacity and splits into roughly equal sizes (a process known as swarming) (Fell et al., 1977). Many obligate cooperators may not be able to join their group of choice, though perhaps the situation is not as restrictive. Regarding assumption 7, though it was introduced for mathematical tractability, the need for collective decision making could introduce a delay in splitting which assumption 7 could approximate (Couzin et al., 2005; Strandburg-Peshkin et al., 2016). Assumption 5 is particularly interesting. Individuals are necessarily limited in information about other groups, but how and to what extent is quite variable. Furthermore, many behaviors evolve such that they appear as if they have more information than possible, i.e. instinct. Overall, we feel the assumptions mimic key features of natural systems while maintaining analytical tractability.

### 4.2. Long-Term Intergroup Dynamics

Over time, the process of groups growing and splitting results in the long-term population dynamics of that species. Under conditions of strong inter-group competition, our model illustrates an initial split followed by both groups simultaneously shrinking to extinction or some non-equilibrial state. This likely manifests as a spatial dynamic. If inter-group competition is spatially dependent, then a group that splits into two, only to remain close, might compete strongly with each, leading to one or both of their extinctions. Rather, on a larger scale, we are likely to see group turnover under conditions of weak inter-group competition. Over longer time scales, our model shows oscillations of the total population size with repeated instances of group extirpations and splitting events (Fig. 5). Similar dynamics can be seen in the reproduction of simple multicellular organisms, a perhaps more advanced form of obligate cooperation (Pichugin et al., 2019).

Despite unstable population dynamics, the overall population size and the number of groups can persist indefinitely within a bounded state. With moderately strong competition, if the groups become unequal in size, then the largest one will grow to maximum size while the smallest is extripated. At this point, the remaining group splits into two and returns to the same original condition. In such a case, though one of the original groups is gone, a new group has taken its place akin to the way a lizard may regrow its tail. With weak competition and group turnover, the overall population exhibits a stable limit cycle. Smaller groups are constantly being extirpated, resetting the system to a previous state. This bounds the range of size over which the overall population will cycle. Old groups are extirpated and new groups are created which function identically to the old groups. In addition, while the population dynamics may be unstable, the distribution of group sizes can be relatively stable.

Studies of long-term population dynamics which focus on the overall population but do not assess the fates of individual groups can attribute fluctuations in population dynamics to external factors. For example, recent population decline to near extirpation of the Isle Royale wolves is attributed to genetic inbreeding or predator-prey dynamics, and a call for human-mediated immigration of new wolves into the Isle Royale population (Hedrik et al. 2014). While there are many environmental factors that may contribute to extirpations, especially those anthropogenic in nature, our model provides support that extirpations and group turnover may simply be an intrinsic property of social animals, a base upon which other factors may be added.

### 4.3. The Limits of Our Model

Altering the assumptions of our model results in different system dynamics, some of them being quite trivial. For example, little happens when we alter the function (eq. 1) for group dynamics while still retaining within-group limits to growth, mostly resulting in the ability to split into multiple groups and a budding dynamic (see SI). Other alterations have larger impacts. Removing the within-group limits to growth removes the behavioral dynamics as individuals will seek to form the grand coalition (a.k.a. the largest group possible) and not leave the group or force out other members. In this grand coalition, fitness is positive, and the group will continually expand. If individuals are sorted into multiple groups and competition forces all group fitness to zero, then the groups are at an unstable population equilibrium.

Removing assumption 6 (individuals cannot join groups) allows for the fusion of smaller groups which can engender some stability (but not full stability) into the system. If these groups can fuse into larger groups, then they can escape extirpation by being in groups larger than the extinction threshold as well as reducing intergroup competition. The inclusion of fusion reduces the upper and lower bounds of the non-equilibrial dynamics associated with total population size and group number (see SI for a specific example). It must be noted though that even with fusion, population dynamics are not truly stable and still result in limit cycles. Removing assumption 7 (having behavioral dynamics at all times, not just at the fitness equilibrium) may create a source-sink dynamic in which individuals are forced out of a group only to subsequently die. This would amount to a behaviorally mediated self-regulation of population size. This only occurs though if assumption 6 also holds; otherwise, should evicted individuals survive, they could form successful groups.

If we alter assumption 5 and let individuals have perfect information, they may not choose to split a group if inter-group competition is strong. At the core of the splitting process are individuals seeking a group size that maximizes fitness. If competition is strong ((*α*_*r*_ ≥ *T*_1_), then it is always better for individuals to be in one group of size *M* than two groups of size *x*_*i*_ and *M* − *x*_*i*_ (0 < *x*_*i*_ < *M*) since the fitness cost of intergroup competition always outweighs the fitness benefits of being inside a group. Group splitting becomes a kind of “mutually assured destruction. If competition is weak (*α*_*r*_ < *T*_1_), then eventually intra-group competition will exceed inter-group competition. Even with perfect information, splitting will still occur, the only difference being that optimal group size is greater than previously 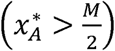.

We can even relax the central tenet of obligate cooperation (assumption 1). Instead, what if there are no fitness benefits to association, only a difference in fitness costs 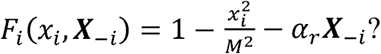 In such a case, individuals seek a group size which minimizes their cost, and there can be a splitting dynamic and unstable population dynamics similar to what we have analyzed. The addition of cooperation guarantees an optimal group size of greater than 1. Under perfect information, there will only be an optimal group size if intergroup competition is not so strong as to create mutually assured destruction, yet not so weak as to have individuals choosing to be by themselves. Under limited information, then no matter the strength of inter-group competition, individuals will choose not to associate with others.

Overall, we feel that our results are robust enough and general enough to apply to a wide variety of systems. Changes to the assumptions may alter the specific dynamics (i.e. extinction versus unstable equilibria versus turnover), but the mismatch between population dynamics and behavioral dynamics remains. Because the general argument for instability relies on this mismatch, obligately cooperative systems should still be unstable.

### 4.4. Conservation Implications

While much has been learned from the social structure of species of high conservation concern (Pusey et al. 2007), our findings suggest future research and conservation efforts should add inter-group dynamics as a major driver for maintaining species population. Our modelling results invite more attention to measuring the size of intra-group Allee effects and the strengths of inter-group competition. Stephens and Sutherland (1999) and Courchamp et al. (2008) focus on the conservation implications of Allee effects in the context of species exploitation, habitat loss, and habitat fragmentation. Long-range dispersal is an important mechanism of species’ range expansions, and dispersal due to group splitting may provide further insights into Allee effects and biological invasions (Lodge, 1993; Taylor and Hastings, 2005).

Our results point to four main effects relevant for the conservation of cooperative species. Firstly, these dynamics are intrinsic and will happen regardless of the environmental conditions. Simply managing for greater environmental stability will not prevent group extirpation or collapse. Persistent, and not necessarily stable, populations should be the goal. Secondly, the average overall population size over time will be smaller than the potential carrying capacity, and the overall population more prone to total extinction due to the constant fluctuations. Therefore, maximizing overall population is critical. Thirdly, there must be a sufficiently large population to allow smaller groups to fuse. The smaller the group, the more likely it is below its internal extinction threshold. By fusing, these smaller groups can avoid the extinction threshold which also has the benefit of boosting average overall population size. And lastly, stronger inter-group competition is more likely to lead to unstable dynamics. Diminished resources and lack of territory between groups can enhance competition and lead to greater instability.

All four of the reasons point to the disproportionate impact that habitat fragmentation and loss should have on obligate cooperative species. Often conservation practices are implemented over smaller scales, with protection for species being implemented in a distinct area of land or for a specific group of that species. For such species, single large conservation areas may be preferable over several small ones (SLOSS) (MacArthur and Wilson 1967, Diamond 1975, Simberloff and Abele 1982). A single large conservation area not only mitigates issues such as inbreeding depression but also may stabilize the population dynamics of obligately cooperative species. Conservation areas should be large enough to harbor multiple groups ensuring minimal inter-group competition. Numerous widely dispersed groups can also withstand environmental stochasticity and may permit a robust fusion process that stabilizes overall population dynamics.

### 4.5. Prospectus

In this paper, we provide a simple model to derive the population dynamics of obligately cooperative species. Our work offers a starting point for further analysis of cooperative species population dynamics to which additional dynamics can be added. From this model, we can develop more realistic models by altering the assumptions of our model or adding features. While we focus on the process of group formation in this study, other important mechanisms for cooperative groups are group maintenance and transformation (Bourke 2011). Such processes would include movement between groups before equilibria, hierarchy and dominance within the group, meta-population and spatial dynamics, source-sink dynamics, evolution, explicit consumer-resource dynamics, and variations on intergroup competition including non-linear competition, asymmetric competition, exploitative vs. interference competition, and fixed and variable intergroup costs. Demographic stochasticity and kin structure are features of smaller obligate cooperative groups not included in this model. Evolution in particular may yet prove fruitful; one way a group may escape the effects of competition in this model is by increasing its growth rate. While the term *α*_*r*_ was used as inter-group competition in this manuscript, it also includes the growth rate. Higher intra-group growth rates concomitantly reduce *α*_*r*_. We can see this in Figure 5. After the second splitting event, the dark red group that is extirpated has a larger maximal group size and larger initial size (and therefore exerts greater competitive force) but a smaller growth rate than either the purple or gold group which both persist. This suggests that a higher growth rate within a group alleviates inter-group competition more so than a larger group size. This aligns with the hypothesis that the evolution of eusociality and division of reproductive work is due to group competition (Reeve and Hölldobler, 2007).

With more complex models of formation, maintenance, and transformation, we can compare the mathematical results against real-world data. We see many examples of group splitting and turnover in nature. Group splitting occurs among rhesus monkeys, lions, sponges, male hyenas, invasive Argentine ants, and orcas (Chepko-Sade and Sade, 1979; Dittus, 1988; Pusey and Packer, 1987; Blanquer et al., 2009; Holekamp et al., 1993; Suarez et al. 2000; Stredulinsky, 2016). We see group turnover among Isle Royale wolves, large primates, wild dogs, elephants, mole-rats, mongooses, spiders, and chimpanzees (Peterson and Page, 1988; Ripple and Beschta 2012; Kalpers et al., 2003; Burrows, 1991; Armbruster and Lande, 1993; Parker and Graham 1989; Jarvis et al., 1994; Aviles, 1997; Clutton-Brock et al., 1999; Goodall 1986). In particular, the Damaraland mole-rats display a process much like the model. Smaller, newly founded groups are more likely to die out due to competition from larger, more established groups (Jarvis et al., 1994). Intergroup competition seems to be a factor in group extirpations of chimpanzees (Goodall 1986). By comparing simulated data from our models to real-world data such as bee swarming or the colonization of wolves in new areas (Oldroyd et al., 1997; Peterson and Page, 1988), we may be able to test how the mismatch between behavioral and population dynamics govern the loss and formation of groups. Because obligately cooperative species often have a significant impact on the ecosystem, whether through ecosystem engineering, their status as keystone species, or accounting for a significant percentage of the biomass of the ecosystem (in some species, all three), it is imperative that ecologists understand the population dynamics of these species (Jones et al., 1994; Ripple and Beschta, 2012; Hoelldolber and Wilson, 1990). Better knowledge will help ecologists and wildlife conservations better manage and save their populations and the ecosystems in which they live (Stephens and Sutherland, 1999).

## Acknowledgements

The authors wish to thank C.J. Whelan, K. Staňková, and G.G. McNickle for their discussion, review, and comments on the paper. AH wishes to thank the NSF for funding his graduate studies. This material is based upon work supported by the National Science Foundation Graduate Research Fellowship under Grant Nos. DGE-0907994 and DGE-1444315. Any opinion, findings, and conclusions or recommendations expressed in this material are those of the author(s) and do not necessarily reflect the views of the National Science Foundation.

---

**Table 1:** A reference table of all the parameters used in this paper.

| Symbol | Definition |
| --- | --- |
| $n_g$ | The number of groups in a population |
| $x$ | The vector of all group sizes |
| $x_i$ | The size of group $i$ |
| $x_{-i}$ | The vector of all group sizes except for group $i$ |
| $X_{-i}$ | The size of the rest of the population |
| $M$ | Maximum potential group size |
| $r$ | A growth rate scaling factor |
| $\alpha$ | The strength of intergroup competition |
| $\alpha_r$ | The ratio between the strength of intergroup competition and the growth rate scaling factor |
| $F(\cdot)$ | The fitness function of a group in our specific model |
| $A(\cdot)$ | The association function of a group in our specific model |
| $x_F^*$ | Equilibrium group size |
| $x_A^*$ | Optimal group size |
| $T_1$ | The main threshold of $\alpha_r$ that divides competition into weak and strong |
| $T_2$ | The secondary threshold of $\alpha_r$ that divides strong competition into moderately strong and extremely strong |
| $\chi^*$ | The smaller root of the fitness function when $x_r > 0$ |
| $\psi^*$ | The larger root of the fitness function when $x_r > 0$ |
| $x_{s,n}$ | The size of the smallest group in a population after the $n$-th split |
| $\chi_{s,n}^*$ | The smaller root of the smallest group's fitness function after the $n$-th split |
| $\psi_{s,n}^*$ | The larger root of the smallest group's fitness function after the $n$-th split |
| $X_{-i,n}$ | The size of the rest of the population from the perspective of the smallest group after the $n$-th split |
| $F_i(\cdot)$ | The fitness function of group $i$ |
| $A_i(\cdot)$ | The association function of group $i$ |
| $x^*$ | A solution to the equation $F_i(x^*) = 0$ |
| $\tilde{x}_i$ | The optimal group size of group $i$ such that $\tilde{x}_i < x_i^*$ |
| $i^+$ | The set of all groups with positive population size |

## Notes

### Competing Interest Statement

The authors have declared no competing interest.

### Summary of Updates

Wording and sections edited to clarify ideas and results

