## Supplementary Information for "Unstable Population Dynamics in Obligate Co-Operators"

**Further Model Analysis**

In this section, we go into the full analysis of our model. Once again, our equation was

$$F(x_{i},x_{R})=r\left( \frac{x_{i}}{M}-\frac{x_{i}^{2}}{M^{2}} \right)-\alpha x_{R}$$

(1)

where $x_{i}$ is the size of group $i$, $r$ is a constant that scales the growth rate, $M$ is maximum group size, $\alpha$ is strength of intergroup competition, and $x_{R}$ is population size of all other groups ($x_{R}=\sum x-x_{i}$). For simpler notation, we divide by $r$ to get

$$F(x_{i},x_{R})=\frac{x_{i}}{M}-\frac{x_{i}^{2}}{M^{2}}-\alpha_{r}x_{R}$$

(2)

where $\alpha_{r}=\frac{\alpha}{r}$, the ratio of intergroup competition strength to growth rate. Since we assume that $r$ is equivalent for all groups, dividing by $r$ does not impact the underlying dynamics in any way.

As stated before, the first two terms govern the intragroup dynamics (benefits of cooperation and costs of competition respectively) while the last term represent intergroup dynamics, specifically competition.

**Equilibria**

In order to find the fitness equilibrium, $x_{F}^{*}$, we set the fitness function equal to 0. Therefore, $F(x_{i},x_{R})=\frac{x_{i}}{M}-\frac{x_{i}^{2}}{M^{2}}-\alpha_{r}x_{R}=0$. Using the quadratic equation, we get

$$x_{F}^{*}=\frac{M}{2}\left( 1\pm\sqrt{1-4\alpha_{r}x_{R}} \right)$$

(3)

Taking the derivative of $F(x_{i},x_{R})$ shows us that the larger of the two equilibria is the stable equilibrium while the smaller one is unstable.

The other important equilibrium is the association equilibrium. To get the association function, we take the derivative of the fitness function with respect to $x_{i}$ holding all other group sizes constant.

$$A(x_{i})=\frac{\partial F\left( x_{i},x_{R} \right)}{\partial x_{i}}=\frac{1}{M}\left( 1-\frac{2x_{i}}{M} \right)$$

(4)

The association equilibrium is determined by setting the association function to 0, resulting in the equilibrium $x_{A}^{*}=\frac{M}{2}$. Taking the derivative of $A(x_{i})$ reveals that it is negative at $x_{A}^{*}$ which confirms it as a maximum and the members’ preffered group size.

As mentioned earlier, $x_{F}^{*}$ and $x_{A}^{*}$ do not match in the absence of intergroup competition, leading to futher dynamics.

**Fitness Dynamics of Two Groups**

Competition between two groups now induces a strong Allee effect with results dependent upon the strength of inter-group competition $\alpha_{r}$. Let us denote the groups’ sizes as $x=(x_{1},x_{2})$ with $x^{*}$ denoting an equilibrium. If $\alpha_{r}\geq\frac{1}{M}$, then there will be three equilibria: $x^{*}=\left( 0,M \right)$, $x^{*}=(M,0)$, and $x^{*}=(0,0)$. The first two equilibria are locally stable; with whichever has the largest group size outcompeting the other. The third equilibrium is saddle point stable – locally unstable except when group sizes are equal.

If $\frac{1}{3M}<\alpha_{r}<\frac{1}{M}$, then there is a new interior equilibrium where $x^{*}=\left( M-\alpha_{r}M^{2},M-\alpha_{r}M^{2} \right)$in addition to the previous 3. This new interior equilibrium now has saddle point stability while the equilibrium of $x^{*}=(0,0)$ is fully unstable. $x^{*}=\left( 0,M \right)$ and $x^{*}=(M,0)$ both remain locally stable.

Lastly, when $0<\alpha_{r}<\frac{1}{3M}$, there are two new interior equilibria: $x^{*}=\left( \frac{M}{2}\left( 1+\sqrt{1-4\alpha_{r}x_{2}} \right),\frac{M}{2}\left( 1-\sqrt{1-4\alpha_{r}x_{1}} \right) \right)$ and $x^{*}=\left( \frac{M}{2}\left( 1-\sqrt{1-4\alpha_{r}x_{2}} \right),\frac{M}{2}\left( 1+\sqrt{1-4\alpha_{r}x_{1}} \right) \right)$. These two new interior equilibria are partially stable while $x^{*}=\left( M-\alpha_{r}M^{2},M-\alpha_{r}M^{2} \right)$now becomes locally stable. In the previous case when $\alpha_{r}>\frac{1}{3M}$, the instability of the equilibria was driven by inter-group competition. Because competition was stronger, any size advantage led the smaller group to be driven to extinction. In this case of weak competition though, the Allee effect and its associated extinction threshold determines the outcome of within and between group population dynamics. If initial group sizes are both above their extinction thresholds, then coexistence will result at the interior equilibrium. If initial group size has one group below its extinction threshold, then it will go extinct and the other group’s size will go to $M$. As intergroup competition goes to 0, the interior equilibrium of $x^{*}=(M,M)$ becomes globally stable and the three equilibria of $x^{*}=(0,0)$, $x^{*}=(M,0)$ and $x^{*}=(0,M)$ become unstable.

Equilibiral and extinction dynamics present themselves when we have two groups of equal size smaller than $x_{A}^{*}=\frac{M}{2}$ at a population equilibium. We can represent this group size as $x_{F}^{*}=cM$, $0\leq c\leq\frac{1}{2}$. Because only two groups, $x_{1}$ and $x_{2}$, exist at the same $x_{F}^{*}$, we get equation 5.

$$F(x_{F}^{*},x_{F}^{*})=\frac{x_{F}^{*}}{M}-\frac{(x_{F}^{*})^{2}}{M^{2}}-\alpha_{r}x_{F}^{*}$$

(5)

Plugging $cM$ into the equation for $x_{F}^{*}$, we get $F(cM,cM)=c-c^{2}-\alpha_{r}cM$. Setting the equation equal to zero and solving for $\alpha_{r}$, we get $\alpha_{r}(c)=\frac{1-c}{M}$. Knowing this, the threshold for equilibrial dynamics $T_{1}$ occurs when $c=\frac{1}{2},\alpha_{r}\left( \frac{1}{2} \right)=\frac{1}{2M}$ and the threshold for extinction dynamics $T_{2}$ occurs when $\alpha_{r}(c)$ is maximized at $c=0,\alpha_{r}(0)=\frac{1}{M}$.

We can further analyze the Jacobian matrix to confirm the instability of these equilibrial states. We write the entire differential equation as $\frac{dx_{i}}{dt}=x_{i}\cdot F(x_{i},x_{j})=\frac{x_{i}^{2}}{M}-\frac{x_{i}^{3}}{M^{2}}-\alpha_{r}x_{i}x_{j}$. Therefore,

$$\frac{\partial\left( \frac{dx_{i}}{dt} \right)}{\partial x_{i}}=\frac{2x_{i}}{M}-\frac{3x_{i}^{2}}{M^{2}}-\alpha_{r}x_{j}$$

(6)

$$\frac{\partial\left( \frac{dx_{i}}{dt} \right)}{\partial x_{j}}=-\alpha_{r}x_{i}$$

(7)

Our Jacobian, $J$, of our two groups, $x_{1}$ & $x_{2}$, now looks like

$$\left[ \begin{matrix} \frac{2x_{1}}{M}-\frac{3x_{1}^{2}}{M^{2}}-\alpha_{r}x_{2} & -\alpha_{r}x_{1} \\ -\alpha_{r}x_{2} & \frac{2x_{2}}{M}-\frac{3x_{2}^{2}}{M^{2}}-\alpha_{r}x_{1} \end{matrix} \right]$$

With $\alpha_{r}=\frac{1-c}{M}$ & $x_{1}=x_{2}=x_{F}^{*}=cM$, we substitute these values into our jacobian to get

$$\left[ \begin{matrix} 2c-3c^{2}-c-c^{2} & -c-c^{2} \\ -c-c^{2} & 2c-3c^{2}-c-c^{2} \end{matrix} \right]$$

This symmetrical matrix gives us the eigenvalues $\lambda_{1}=c\left( 2-3c \right), \lambda_{2}=-c^{2}$. Since $-c^{2}\leq0$ always, we look to $\lambda_{1}$ to understand the dynamics. If the two groups have positive sizes at equilibrium, then $0<c\leq\frac{1}{2}$; therefore $\lambda_{1}>0$ meaning instability. $\lambda_{1}$ is only negative when $c>\frac{2}{3}$, $\alpha_{r}<\frac{1}{3M}$. If both groups are extinct $x_{F}^{*}=0$, we get an all $0$ matrix and $\lambda_{1}=\lambda_{2}=0$.

**Fitness Dynamics of More than Two Groups**

We can use the same analysis from the previous section to analyze the fitness dynamics of a system with more than two groups. Once again, we look only at the interior equilibrium where all individuals have the same group size to understand thresholds. That is because there are only four types of equilibria:

1. all $x_{i}=0$
2. one group at size $M$ and the rest at $0$
3. some at the smaller root and some at the larger root
4. and all at the same root whether larger or smaller

The first equilibrium is trivial, the second equilibrium is a special case of the fourth equilibrium where there is only one group with positive size, and the third equilibrium is always (partially) unstable due to the extinction threshold. Therefore, the last equilibrium is key to the dynamics.

We start by looking at strength of $\alpha_{r}$. We write the fitness dynamics of a single group as

$$F(x_{F}^{*},x_{F}^{*})=\frac{x_{F}^{*}}{M}-\frac{(x_{F}^{*})^{2}}{M^{2}}-(n_{g}-1)\alpha_{r}x_{F}^{*}$$

(8)

Here, $n_{g}$ is the number of groups in our system at the equilibrium $x_{F}^{*}$. Once again setting $x_{M}^{*}=cM$ and solving for $\alpha_{r}$, we get $\alpha_{r}(c)=\frac{1-c}{(n_{g}-1)M}$. As the number of groups increases, $\alpha_{r}$ must decrease for there to be an interior equilibrium.

We can also once again use Jacobian analysis to get at the shift our equilibrium’s stability. The Jacobian turns out to be of the form

$$\left[ \begin{matrix} b+a & a & a & \ldots& a \\ a & b+a & a & \ldots& a \\ a & a & b+a & \ldots& a \\ \ldots& \ldots& \ldots& \ldots& \ldots\\ a & a & a & \ldots& b+a \end{matrix} \right]$$

where $a=-c-c^{2}$ and $b=2c-3c^{2}$. We can say $J=A+B$ where $A$ is a $n_{g}$x$n_{g}$ matrix with all the values equal to $a$ and $B$ is the identity matrix of size $n_{g}$ multiplied by the scalar $b$, $B=bI_{n_{g}}$. Adding a scaled identity matrix to another matrix merely shifts the eigenvalues by the scalar. Therefore, the eignevalues of $J$ are the eigenvalues of $A$ plus $b$. The eigenvalues of a matrix where all elements are the same are one eigenvalue equal to the size of the matrix times the element with the rest being $0$. Therefore, the eigenvalues of $A$ are one eigenvalue equal to $n_{g}a$ and the rest $0$. As such, the eigenvalues of $J$ are one eigenvalue of $n_{g}a+b$ and the rest equal to $b$.

Solving for when these eigenvalues are less than 0 yields the inequalities $c>\frac{2-k}{3+k}$ and $c>\frac{2}{3}$. This is functionally the same as the two-group system where $x_{F}^{*}>\frac{2}{3M}$ for this interior equilibrium to be stable. The only difference is that $\alpha_{r}$ is scaled by $\frac{1}{n_{g}-1}$.

**Group Turnover**

When $T_{1}>\alpha_{r}>0$, the eventual outcome is group turnover and localized extinction of groups. This only occurs though after a buildup in the population. Therefore, this dynamic is dependent on $x_{R}$. The size of the new groups formed are always determined by the value of the largest root $\frac{M}{2}\left( 1+\sqrt{1-4\alpha_{r}x_{R}} \right)$ minus $x_{A}^{*}$. If the size of these new groups are smaller than extinction threshold $\frac{M}{2}\left( \sqrt{1-4\alpha_{r}x_{R}} \right)$, then they will automatically go extinct. This creates an upper bound on the size on the rest of the population when group turnover begins. It must be remembered though that the new, smaller group was combined with one of the larger groups before splitting. Therefore we replace $x_{R}$ with $x_{R}'=x_{R}-\frac{M}{2}$ and the size of the new groups are $\frac{M}{2}\left( \sqrt{1-4\alpha_{r}x_{R}'} \right)$. Now to solve for the $x_{R}$ threshold, this value must be equal to the value of the extinction threshold. This leads to $\frac{M}{2}\left( \sqrt{1-4\alpha_{r}x_{R}'} \right)=\frac{M}{2}\left( 1-\sqrt{1-4\alpha_{r}x_{R}} \right)$. Solving through for $x_{R}$, we get

$$x_{R}=\frac{1}{4\alpha_{r}}-\frac{1}{16\alpha_{r}}(1-2\alpha_{r}M)^{2}$$

(9)

With the value of $x_{R}$ needed to start turnover, the numbers of groups needed, $n_{g}$, can be figured out. The total population is the $x_{R}+\frac{M}{2}\left( \sqrt{1-4\alpha_{r}x_{R}} \right)$. This can be divided by the large root of the fitness function $\frac{M}{2}\left( 1+\sqrt{1-4\alpha_{r}x_{R}} \right)$ to find the minimum number of groups needed to start turnover. Because of symmetry, the groups grow by a doubling process; the number of groups after the $n$-th split is the initial number of groups, $n_{g_{0}}$, times $2^{n}$. Extirpations will occur with group numbers such that $n_{g_{0}}\cdot2^{n}<n_{g}<n_{g_{0}}\cdot2^{n+1}$.
While we feel this model is certainly interesting, we recognize that it does not capture the full variation of group dynamics. We can generalize the function to get more variety in the relationship between group size and fitness.

**Generalized Allee**

We can now generalize the "cooperative" and "competitive" intra-group components of the equations by raising them to exponents to get the equation:

$$F(x_{i},x_{R})=\left( \frac{x_{i}}{M} \right)^{l}-\left( \frac{x_{i}}{M} \right)^{l+m}-\alpha_{\hat{r}}x_{R}$$

(10)

with $l,m>0$. Because the costs of competition must outstrip the benefits of cooperation at some point for there to be a positive fitness equilibrium, the exponent on the second term must necessarily be greater than the exponent on the first term. We can rewrite the equation as

$$F(x_{i},x_{R})=\left( \frac{x_{i}}{M} \right)^{l}\left( 1-\frac{x_{i}}{M} \right)^{m}-\alpha_{\hat{r}}x_{R}$$

(11)

In the previous section, we were able to fully describe the dynamics since $l=m=1$. Allowing $l$ and $m$ to be any combination of positive numbers is quite more interesting but harder to describe the dynamics. We can make an attempt at describing some interesting properties when the function is generalized.

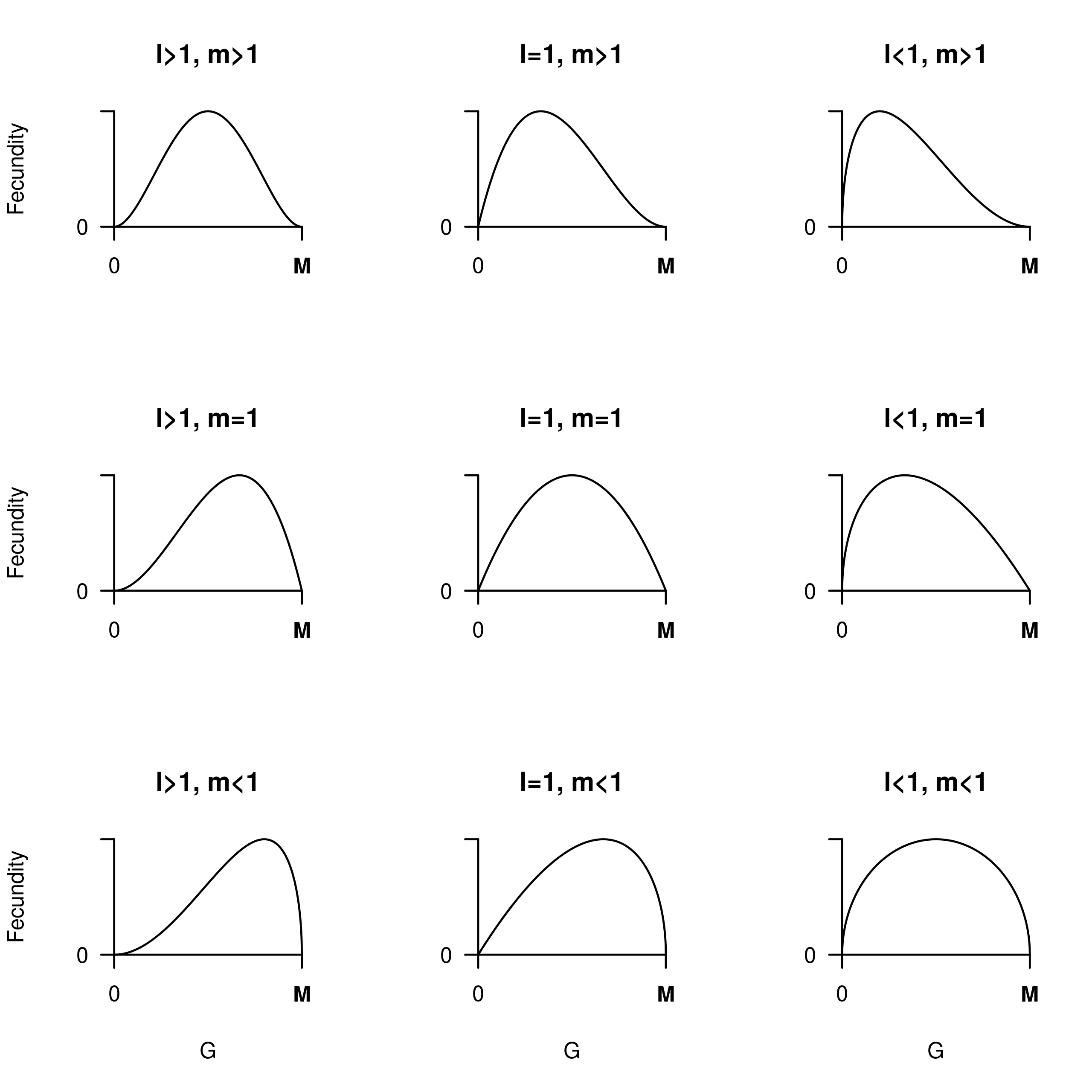

SI Figure 1. All possible combinations of l, m >, =, < 1

There are four things to note about this equation. Firstly, as $l$ and $m$ changes, $F(x_{A}^{*},x_{R})$ changes with it. In this case, we introduce $\hat{r}$ which scales appropriately to give reasonable dynamic speeds. One possibility is to say $\hat{r}$ is on the order of $r\cdot2^{l+m-2}$. Secondly, an even or non-integer $m$ leads to strange dynamics beyond $M$. If $m$ is even, the fitness function is a local minimum at $M$, rising to infinity after $M$. If $m$ is not an integer, then values beyond $M$ yields complex numbers with 0 real parts. Since the groups compete with each other, only the fitness function between 0 and $M$ really matters. Therefore, we can ignore group sizes greater than $M$ and assume it is monotonically negative beyond $M$. Thirdly, while we cannot find a general solution to $x_{F}^{*}$, we get the general solution $x_{A}^{*}=\frac{lM}{l+m}$. And lastly, when $l,m\neq1$, the shape of the function changes dependent on the size of the exponents and which exponents change. With an exponent greater than 1, the function kinks inward; outward when less than 1. As well, changing $l$ causes the kinking to the left of $x_{A}^{*}$ while changing $m$ causes kinking to the right (SI Fig. 1). This has implications for the dynamics, namely both $\alpha_{\hat{r}}$ and $x_{R}$ now act non-linearly on the roots function.

With this information, we can divide the potential dynamics into two broad types based on what happens when: $l\geq m$ and $l<m$. In the first case, when starting from a single group at $x_{F}^{*}$ will split into two, one of size $\frac{lM}{l+m}$ and $\frac{mM}{l+m}$. In the second case, we could get multiple groups depending upon the ratio between $l$ and $m$ (which will be explained later). We look at the case when $l\geq m$ first.

$\boldsymbol{l\geq m}$

With $l\geq m$, a group at $x_{F}^{*}$ greater than $x_{A}^{*}$ will split into two. We will call these groups $x_{1}$ of size $x_{1}=x_{A}^{*}=\frac{lM}{l+m}$ and $x_{2}$ of size $x_{2}=\frac{mM}{l+m}$. Let $F_{1}=F(x_{1},x_{2})$ and $F_{2}=F(x_{2},x_{1})$. We can characterize the eventual dynamics based on the fact whether each group persists or goes extinct in the resulting dynamics after splitting. If both groups eventually go extinct, we call that an extinction dynamic; if one group persists while the other goes extinct, we call that a budding dynamic; if both groups eventually persist with initial fitness less than or equal to 0, we call this equilibrial dynamics; and if both groups eventually persist having initial fitness greater than 0, we can get many dynamics including group turnover and budding.

Now, we can then create a framework based upon whether the fitness of these groups are greater than, equal to, or less than 0.

|  | $F_{2}>0$ | $F_{2}=0$ | $F_{2}<0$ |
| --- | --- | --- | --- |
| $F_{1}>0$ | Turnover,  Budding,  Other | Budding | Budding |
| $F_{1}=0$ | N/A | Equilibrial | Budding |
| $F_{1}<0$ | N/A | N/A | Equilibrial,  Extinction,  Budding |

Table 1. Possible dynamics between two cooperative groups when $l\geq m$. $F_{1}$ and $F_{2}$ represent the fitness of each group

With this table, we can now begin to analyze each possibility. First thing to notice is that the lower triangle of the table is impossible. This is because $F_{2}$ cannot be greater than $F_{1}$. The fitness function rises monotonically from 0 to $x_{A}^{*}$. Since $l\geq m$, $x_{1}=x_{A}^{*}\geq x_{2}$ making $F_{1}\geq F_{2}$. This yields six possibilites, three of which will be shown to be the same, leaving four to be analysed below.

$\boldsymbol{F}_{\boldsymbol{1}}\boldsymbol{=}\boldsymbol{F}_{\boldsymbol{2}}\boldsymbol{=0}$

If $F_{1}=F_{2}=0$, then we know both groups will persist in a partially stable equilibrium. In fact, we can solve for some necessary conditions. We can begin by solving for $F_{1}$ and setting it to 0 obtaining $F_{1}=\left( \frac{x_{1}}{M} \right)^{l}\left( 1-\frac{x_{1}}{M} \right)^{m}-\alpha_{\hat{r}}x_{2} = \left( {\frac{lM}{l+m}}/M \right)^{l}\left( 1-{\frac{lM}{l+m}}/M \right)^{m}-\alpha_{\hat{r}}\frac{mM}{l+m} = 0$. Therefore, $\left( \frac{l}{l+m} \right)^{l}\left( \frac{m}{l+m} \right)^{m}=\alpha_{\hat{r}}\frac{mM}{l+m}$ yielding $\left( \frac{l}{l+m} \right)^{l}\left( \frac{m}{l+m} \right)^{m-1}\frac{1}{M}=\alpha_{\hat{r}}$. Similarly solving for $F_{2}$ gives us $\left( \frac{m}{l+m} \right)^{l}\left( \frac{l}{l+m} \right)^{m-1}\frac{1}{M}=\alpha_{\hat{r}}$. We can now say

$$\left( \frac{l}{l+m} \right)^{l}\left( \frac{m}{l+m} \right)^{m-1}\frac{1}{M}=\alpha_{\hat{r}}=\left( \frac{m}{l+m} \right)^{l}\left( \frac{l}{l+m} \right)^{m-1}\frac{1}{M}$$

(12)

Reducing this equation, we arrive at the fact that $l=m$. This is a necessary but not sufficient condition for a partially stable equilibrium. In order to have $F_{1}=F_{2}=0$, $\alpha_{\hat{r}}$ must also equal to $\frac{1}{2^{2m-1}M}$. This is because the condition $l=m$ only guarantees that $F_{1}=F_{2}$, not whether they are positive, negative, or zero. In fact, the only way to get $F_{1}=F_{2}$ is to have $l=m$. Writing the full equation out, we get $\frac{l^{l}m^{m}}{(l+m)^{l+m}}-\alpha_{\hat{r}}\frac{m}{l+m}M=\frac{m^{l}l^{m}}{(l+m)^{l+m}}-\alpha_{\hat{r}}\frac{l}{l+m}M$. Solving through, we get

$$\frac{(l^{l-m}-m^{l-m})(ml)^{m}}{(l+m)^{l+m-1}}=(m-l)\alpha_{\hat{r}}M$$

(13)

If $l>m$, then the RHS is negative while the LHS is positive; vice versa when $m<l$. This leads to a proof by contradiction. Only when $l=m$ does the RHS equal the LHS. More generally, if $l=m$ and $\alpha_{\hat{r}}>\frac{1}{2^{2m-1}M}$, then $F_{1}=F_{2}<0$; similarily, if $\alpha_{\hat{r}}<\frac{1}{2^{2m-1}M}$, then $F_{1}=F_{2}>0$. We will explore more about $l=m$ in a later section.

$\boldsymbol{F}_{\boldsymbol{1}}\boldsymbol{>}\boldsymbol{F}_{\boldsymbol{2}}\boldsymbol{;}\boldsymbol{F}_{\boldsymbol{1}}\boldsymbol{>}\boldsymbol{F}_{\boldsymbol{2}}\boldsymbol{=0;}\boldsymbol{F}_{\boldsymbol{1}}\boldsymbol{=0>}\boldsymbol{F}_{\boldsymbol{2}}$

With these 3 possibilities, we get a new phenomenon in which one group ultimately persists while the other goes extinct, hereafter referred to as budding. With budding, after the group splits, $x_{2}$ goes extinct while $x_{1}$ grows back to $M$. At this point, $x_{1}$ splits in the same way as before with new group going extinct and $x_{1}$ growing again; it is quite similar to the turnover dynamics only with a single group instead of multiple groups.

Budding dynamics are quite natural to see when $F_{1}>0>F_{2}$. When $F_{1}>0=F_{2}$ or $F_{1}=0>F_{2}$, we eventually get to the state where $F_{1}>0>F_{2}$. In the first case, since $F_{1}>0$, $x_{1}$ will grow and exert greater inter-group competition on $x_{2}$ forcing $F_{2}$ below zero. At this point, $F_{1}>0>F_{2}$. In the second case, since $F_{2}<0$, $x_{2}$ will shrink and exert less inter-group competition on $x_{1}$ boosting $F_{1}$ above zero at which point we get $F_{1}>0>F_{2}$. Therefore, all three states are functionally equivalent.

To solve for conditions for these three possibilities, we modify the equality sign in equation 12 to fit the scenario accordingly. This means we arrive at

$$\left( \frac{l}{l+m} \right)^{l}\left( \frac{m}{l+m} \right)^{m-1}\frac{1}{M}>\alpha_{\hat{r}}>\left( \frac{m}{l+m} \right)^{l}\left( \frac{l}{l+m} \right)^{m-1}\frac{1}{M}$$

(14)

or

$$\left( \frac{l}{l+m} \right)^{l}\left( \frac{m}{l+m} \right)^{m-1}\frac{1}{M}>\alpha_{\hat{r}}=\left( \frac{m}{l+m} \right)^{l}\left( \frac{l}{l+m} \right)^{m-1}\frac{1}{M}$$

(15)

or

$$\left( \frac{l}{l+m} \right)^{l}\left( \frac{m}{l+m} \right)^{m-1}\frac{1}{M}=\alpha_{\hat{r}}\left( \frac{m}{l+m} \right)^{l}\left( \frac{l}{l+m} \right)^{m-1}\frac{1}{M}$$

(16)

Regardless, the equation reduces to $l>m$. This is a necessary but not sufficient condition to get budding. In a later section, we show that we cannot get budding dynamics when $l=m$ (proof by contradiction), yet we also show that dynamics other than budding can occur when $l>m$, $F_{1}>F_{2}$.

$\boldsymbol{F}_{\boldsymbol{1}}\boldsymbol{,}\boldsymbol{F}_{\boldsymbol{2}}\boldsymbol{<0}$

If we have $F_{1},F_{2}<0$, then there are three potential dynamics: extinction, equilibrial, and budding. The first two occur when $l=m$ and the third occurs when $l>m$. In fact, when $l>m$ and $F_{1},F_{2}<0$, only budding dynamics will occur which we can show with rough logic. If $l>m$, then $x_{1}>x_{2}$, $F_{1}>F_{2}$. Since $F_{1},F_{2}<0$, both groups will shrink in size, heading towards 0. Since both groups have the same fitness function, $x_{2}$ will always be smaller than $x_{1}$. This means that $F_{1}>F_{2}$ over all time. As $x_{1}$ and $x_{2}$ shrink, $F_{1}$ and $F_{2}$ increase, eventually reaching a point in time where $F_{1}=0>F_{2}$. We can prove this by assuming there exists a point in time where $x_{2}$ is extinct, $x_{2}=0$. At this point, $x_{1}$ faces no competition. With our fitness function, any value in the interval $(0,M)$ gives positive fitness. Since $x_{1}>x_{2}=0$, then $F_{1}>0$. Since we started with $F_{1}<0$, at some point before $F_{1}$ must have crossed the x-axis giving us $F_{1}=0>F_{2}$. Therefore, if $l>m$ and $F_{1},F_{2}<0$, we can only get budding dynamics. As well, when $l=m$, $F_{1}=F_{2}<0$ giving us only equilibrial or extinction dynamics, described in a later section.

$\boldsymbol{F}_{\boldsymbol{1}}\boldsymbol{,}\boldsymbol{F}_{\boldsymbol{2}}\boldsymbol{>0}$

With $F_{1},F_{2}>0$, we can get many dynamics! To understand what happens, we first look at two extremes. The first extreme occurs when $F_{1}=F_{2}$. This happens when $l=m$ as determined before and leads to group turnover. Once again, we put that aside for a later section. The other extreme occurs when we have $l>m$, $x_{1}>x_{2}$, and $\left. F_{2}=\epsilon\downarrow0 \right.$. At this extreme, the dynamics are essentially the same when $F_{1}>F_{2}=0$, giving us budding. This means there is some combination of $l$, $m$, $\alpha_{\hat{r}}$, and $M$ that transitions between turnover and budding. Determining that threshold is quite tricky since it depends upon the both $F_{1}$ and $F_{2}$ as well as the slope at those points. The only way to characterize the dynamics is through the final state of the two groups, specifically the ultimate result of $x_{2}$. As $x_{1}$ grows, it will always end up at an equilibrium greater than $x_{A}^{*}$. Let’s denote the new fecundities when $x_{1}$ first reaches this equilibrium as $F_{1}'$ and $F_{2}'$. At this point, $F_{2}'$ can either be positive, negative, or zero.

If $F_{2}'>0$, $x_{2}$ will keep growing forcing $F_{1}'$ negative due to increased inter-group competition and causing $x_{1}$ to shrink. Eventually, both groups will converge on the same group size. It is hard to determine this size in relation to $x_{A}^{*}$. If it is less than $x_{A}^{*}$, then both with rest at an unstable equilibrial state. If greater than $x_{A}^{*}$, then we have turnover dynamics. We suspect that it is always less than $x_{A}^{*}$ always, but we cannot be certain.

If $F_{2}'<0$, then we have budding dynamics.

If $F_{2}'=0$, $x_{2}$ must be at the smaller root since $x_{1}$ will be at the larger root and they are not equal. Since $x_{1}$ is at a competitive equilibrium, it will split to grant another group $x_{3}$. In this case, $x_{1}=x_{A}^{*}>x_{2}>x_{3}$. We can confidently state that $F_{1}>F_{2}>F_{3}$ since the respective groups are bigger and are ordered in terms of descending competitive effect ($x_{R}$). Since $x_{R}$ for $F_{2}$ did not change, $F_{1}>F_{2}=0>F_{3}$; therefore, $x_{3}$ will always go extinct and $x_{1}$ will always grow. What happens to $x_{2}$ is less certain. If $x_{1}$ grows to its previous state at the same rate that $x_{3}$ goes extinct, then $F_{2}$ will always equal $0$ and the dynamics will simply repeat. If not, then $x_{2}$ will fluctuate and either increase to create the scenario when $F_{2}'>0$, decrease to create the scenario when $F_{2}'<0$, or end up back at the same place.

Fully described dynamics when $l>m$ is hard to come by. What we can say is that if at least $F_{2}\leq0$ immediately after splitting, then we will get budding dynamics. Beyond that, we cannot make any general conclusions. We are able to fully understand the dynamics when $l=m$ regardless of whether $F_{i}>,<,=0$ as seen in the next section.

$\boldsymbol{l=m}$

The only time we can obtain a general solution to $x_{F}^{*}$ is when $l=m$. With $l=m$, we get equation 17

$$F(x_{i},x_{R})=\left( \frac{x_{i}}{M}\left( 1-\frac{x_{i}}{M} \right) \right)^{m} -\alpha_{\hat{r}}x_{R}=0$$

(17)

which, when $F(x_{i},x_{R})=0$, can be transformed into $\frac{x_{i}}{M}\left( 1-\frac{x_{i}}{M} \right)=\left( \alpha_{\hat{r}}x_{R} \right)^{\frac{1}{m}}$.

At this point, the function is essentially identical to the example in Section 2, only with the effects of $\alpha_{\hat{r}}x_{R}$ being non-linear. The roots are of the same form as the previous roots, only now $x_{F}^{*}=\frac{M}{2}\left( 1\pm\sqrt{1-4 \left( \alpha_{\hat{r}} x_{R} \right)^{\frac{1}{m}}} \right)$. Because $l=m$, $F(x_{i},x_{R})$ is also symmetrical with respect to $x_{i}$ over the interval [0,$M$] and $x_{A}^{*}=\frac{M}{2}$.

Now that we have the roots, we can solve for equilibrial dynamics. Using the same $x_{F}^{*}=cM$, we can solve for $\alpha_{\hat{r}}(c)=\frac{c^{m-1}(1-c)^{m}}{M}$; therefore at $T_{1}$, $\alpha_{\hat{r}}\left( \frac{1}{2} \right)=\frac{1}{2^{2m-1}M}$. $T_{2}$ provides a much more interesting case. As we can see, the general formula for $\alpha_{\hat{r}}(c)$ is non-linear. This leads to two separate phenomena depending upon whether $m>1$ or $m<1$.

If $m>1$, then $\alpha_{\hat{r}}(c)$ starts at the value $0$ for $c=0$, rises to a maximum value at $c_{max}$, then falls to the value of $0$ at $c=1$. The value of $\alpha_{\hat{r}}$ at $c_{max}$ is the new threshold value $T_{2}$. We can solve for $T_{2}$ by taking the derivative of $\alpha_{\hat{r}}(c)$ with respect to $c$. Rewriting $\frac{c^{m-1}(1-c)^{m}}{M}$ as $\frac{(c-c^{2})^{m-1}(1-c)}{M}$, we get

$$\frac{\partial\alpha_{\hat{r}}(c)}{\partial c}=\left( \frac{1}{M} \right)\left( \left( m-1 \right)\left( c-c^{2} \right)^{m-2}\left( 1-c \right)\left( 1-2c \right)-\left( c-c^{2} \right)^{m-1} \right)$$

(18)

Setting this equation to $0$, we can get $c_{max}$. Though it may seem hard given that we are potentially dealing with a higher order polynomial, we can rewrite the partial derivative as

$$\frac{\partial\alpha_{\hat{r}}(c)}{\partial c}=\left( \frac{1}{M} \right)(c-c^{2})^{m-2}\left( 1-c \right)\left( \left( m-1 \right)\left( 1-2c \right)-c \right)$$

(19)

Dividing by $\frac{1}{M}(c-c^{2})^{m-2}(1-c)$, we are now left with only the equation $0=(m-1)(1-2c)-c$. Solving for the equation, we get $c_{max}=\frac{m-1}{2m-1}$. Plugging $c_{max}$ into $\alpha_{\hat{r}}(c)$, we get $T_{2}=\frac{1}{M}\frac{(m-1)^{m-1}(m)^{m}}{(2m-1)^{2m-1}}$. The non-linearities of $\alpha_{\hat{r}}(c)$ also restrict the range of potential equilibrial values. A value of $\alpha_{\hat{r}}(c)$ can potentially lead to two equilibrium values $c_{1}M$ and $c_{2}M$, $c_{1}<c_{max}<c_{2}\leq\frac{1}{2}$. Since the populations start off at equals size $x_{A}^{*}$ in order to get equilibrial dynamics, this means that the dynamics will only go to the value $c_{2}M$; any equilibrium of $cM$ where $0<c<c_{max}$ is unobtainable. This means that $c$ now ranges on the interval $\left[ \frac{m-1}{2m-1},\frac{1}{2} \right]$.

When $m<1$, this leads to a strange case. Looking at $\alpha_{\hat{r}}(c)$ as a whole, we see that when $m<1$ it starts at infinity then declines monotonically to $\frac{1}{2^{2m-1}M}$. In order to achieve extintion dynamics when $m<1$, competition must be infinitely strong, making extinction unachievable.

We now look to the Jacobian to understand the stability of the equilibrium values. The full $\frac{dx_{i}}{dt}$ is $x_{i}\left( \left( \frac{x_{i}}{M}-\frac{x_{i}^{2}}{M^{2}} \right)^{m}-\alpha_{r}x_{j} \right)$. The partial with respect to $x_{i}$ is

$$\frac{\partial\left( \frac{dx_{i}}{dt} \right)}{\partial x_{i}}=\left( \frac{x_{i}}{M}-\frac{x_{i}^{2}}{M^{2}} \right)^{m-1}\left( \left( \frac{x_{i}}{M}-\frac{x_{i}^{2}}{M^{2}} \right)+mx_{i}\left( \frac{1}{M}-\frac{2x_{i}}{M} \right) \right)-\alpha_{\hat{r}}x_{j}$$

(20)

and the partial with respect to $x_{j}$ is the same

$$\frac{\partial\left( \frac{dx_{i}}{dt} \right)}{\partial x_{j}}=-\alpha_{r}x_{i}$$

(21)

Our Jacobian, $J$, of our two groups, $x_{1},x_{2}=cM$, now looks like

$$\left[ \begin{matrix} b-\alpha_{\hat{r}}cM & -\alpha_{\hat{r}}cM \\ -\alpha_{\hat{r}}cM & b-\alpha_{\hat{r}}cM \end{matrix} \right]$$

where $b=(c-c^{2})^{m-1}((m+1)c-(2m+1)c^{2})$. Substituting solving for the eigenvalues, we get $\lambda=b-\alpha_{\hat{r}}cM\pm\alpha_{\hat{r}}cM$. Plugging the general solution of $\alpha_{\hat{r}}$ into the eigenvalues, $\lambda_{1}=b,\lambda_{2}=b-2(c-c^{2})^{m}$. $b>0$ for all values of $c\in\left[ \frac{m-1}{2m-1},\frac{1}{2} \right]$ if $m>1$ and $c\in\left( 0,\frac{1}{2} \right]$ if $m<1$. Therefore $\lambda_{1}>0$ for all potential equilibrium values, meaning all potential equilibrium values are unstable. If both groups are extinct, then $\lambda_{1}=\lambda_{2}=0$.

Even though we can solve for $x_{F}^{*}$, we cannot solve for the $x_{R}$ needed for turnover dynamics because of the non-linearities present. Yet, we can still understand the basic dynamics. The roots are essentially the same as the previous example, only with the $m$-th root of the competition part ($\alpha_{r}x_{R}$). Using equation 17, $F(x_{A}^{*},0)={0.25}^{m}$; therefore $\alpha_{\hat{r}}x_{R}<{0.25}^{m}<1$. If $m>1$, then $\left( \alpha_{\hat{r}}x_{R} \right)^{\frac{1}{m}}>\alpha_{\hat{r}} x_{R}$, meaning that $x_{R}$ has a stronger competitive effect on a group compared to when $m=1$ especially at a lower group size. Therefore, group turnover will occur at a lower $x_{R}$. This can be confirmed using graphs (Figure 1). Because of the inward kinks, the roots change very quickly at low competition before becoming more resilient, confirming the need for a smaller $x_{R}$ to get group turnover. The reverse is true when $m<1$. $\left( \alpha_{\hat{r}}x_{R} \right)^{\frac{1}{m}}<\alpha_{\hat{r}} x_{R}$ meaning $x_{R}$ has a weaker competitive effect on group compared to when $m=1$ and a higher $x_{R}$ for turnover to occur. The graphs with their outward kinks confirm this resilience at smaller population sizes.

With $l\geq m$ completed, we can now look at what happens when $l<m$.

$\boldsymbol{l<m}$

If $l<m$, then the function rises quickly on average before falling slowly on average. This leads to an $x_{A}^{*}=\frac{lM}{l+m}<\frac{M}{2}$. Therefore, the single group can split into more than one group depending on the ratio $\frac{l}{l+m}$, specifically $\left\lceil\frac{l+m}{l} \right\rceil$. Determining the dynamics with multiple groups becomes harder though some general results can appear. The key to understanding the system is that in some instances there may be a straggler group. This occurs when $\frac{l+m}{l}$ is not an integer. In this case, there are $n_{g}-1=\left\lfloor\frac{l+m}{l} \right\rfloor$ groups of size $x_{A}^{*}$ and one group of size $M-(n_{g}-1)\cdot x_{A}^{*}$. The dynamics of this group will then go on to determine the dynamics of the entire population.

**Equilibrial and Extinction Dynamics**

In the case of equilibrial and extinction dynamics, we can get general results for the system. Since all groups have fitness less than $0$, all groups will initially decrease. If there is a straggler group, it will go extinct first because it is smallest. At that point, there at $n_{g}-1$ identical groups which will either continue to go extinct or reach some equilibrial state. In that case, we can solve for $T_{1}$ and $T_{2}$. Using the equilibrium $x_{F}^{*}=cM$, we get the equation $\alpha_{r}(c)=\frac{c^{l-1}(1-c)^{m}}{(n_{g}-2)M}$. We can see that the equation is of similar form as when $l=m$, only multiplied by the constant $\frac{1}{n_{g}-2}$. The dynamics are also quite similar: if $l=1$, we get a monotonic decrease from a positive point; if $l<1$, then we get a monotonic decrease but from infinity; and if $l>1$, then our equation rises and then falls from $0$ to $\frac{1}{2}$. This also gives us a general equation for $\alpha_{r}(c)$ which can recapitulate all other specific dynamics.

**Turnover Dynamics**

Group turnover cannot be generally solved for as they require a general picture of the roots of the fitness equation which are unsolveable. We can say though that the dynamics are very similar to when $l\geq m$. Let $F$ be the fitness of all groups at $x_{A}^{*}$ and $F_{s}$ be the fitness of the straggler group. We will get extirpation dynamics when $F>0$. If $F_{s}\leq0$, then the straggler group will go extinct leading to simple, repeatable extirpation dynamics. If $F_{s}>0$, then we get a scenario similar to when $l>m$, $F_{1},F_{2}>0$.

**Stabilizing Influences**

We now expand on the potential list of stabilizing influences from the paper. We take a deeper look at the main stabilizing influence, the fusion process, before delving into more technical and less biologically plausible possibilities.

**Fusion**

In our model, we assumed that groups can only fission; that groups could not join to form larger groups based on their interest. We know that in many cooperative species, smaller groups will fuse to form larger groups. This ability to fuse changes the turnover dynamics. group turnover, each new group’s population is $\frac{M}{2}\left( \sqrt{1-4\alpha_{r}x_{R}} \right)$. Because $\frac{M}{2}\left( \sqrt{1-4\alpha_{r}x_{R}} \right)<\frac{M}{2}$ with positive $\alpha_{R}x_{R}$, new groups will choose to aggregate with each other to form groups of size $\frac{M}{2}$. The most obvious result of this that groups below extinction threshold can bond and survive extinction. This could lead to a higher number of groups before cycling. This is much harder to solve for though due to the complicated dynamics. One solution easier to solve for though is the ability to achieve partially unstable equilibira. If $\frac{M}{2}\left( \sqrt{1-4\alpha_{r}x_{R}} \right)$ multiplied by the number of groups is equal to a multiple of $\frac{M}{2}$, then these groups will bond to form that multiple of groups at $\frac{M}{2}$. This allows for a greater number of half-stable equilibria to be present in the solution.

As an example, if two groups both reach a size $\frac{3M}{4}$, they will split to form 4 groups: two of size $\frac{M}{2}$ and two of size $\frac{M}{4}$. With fusion, the two smaller groups can form a third group of size $\frac{M}{2}$, resulting in a partially unstable equilibrium. If fusion weren’t possible, the smaller groups would have gone extinct instead, resulting in group turnover.

**Other Stabilizing Influences**

We hypothesize three more stabilizing influences. Because these are more technical and not biologically based, we only briefly discuss them.

1) There may be a limit to splitting. We assumed that there was always sufficient resources for the creation of a new group. A limited environmental and resource space may constrain total population size and therefore the number of groups. Each split may reduce optimal group size until it is so small that groups cannot split anymore (optimal group size is 1). In this case though, the system has just recapitulated regular competition between individuals within a population. As well, limited environmental space likely leads to more turnover, not less.
2) Another possibility is that all groups are at maximum fitness and neutrally stable. In this case, minor disturbances will not cause groups to continue to grow or shrink. Such an outcome requires very specific fitness functions likely not to be seen in nature.
3) The last possibility is when $x_{A}^{*}\approx x_{F}^{*}$. In this case, optimal group size is essentially equivalent to maximal group size. We call this the House of Cards scenario because individuals continually add to fitness until a small number of individuals cause fitness to crash to zero. As $x_{A}^{*}$ approaches $x_{F}^{*}$, then the size of the new groups $x_{F}^{*}-x_{A}^{*}$ approaches 0. This though can only exist in a continuous system; $x_{A}^{*}$ will always be different from $x_{F}^{*}$ in a discrete system and like neutral stability requires specific functions likely not seen in nature.
4) Perfect information could create a situation similar to Mutually Assured Destruction. If competition is strong enough, then members may not choose to split the group after reaching the fitness equilibrium as it will lead to less fitness than if they were in a single group. We can write an equation of perfect information as such

$$F\left( x_{i},x_{R} \right)=\frac{x_{i}}{M}-\frac{x_{i}^{2}}{M^{2}}-\alpha_{r}\left( M-x_{i} \right)-\alpha_{r}x_{R}$$

(22)

Here, individuals of group size $M$ could choose to split the group into two new groups: one of size $x_{i}$ and one of size $M-x_{i}$. Equation 22 gives the fitness of the first group. If $x_{i}$ is entirely negative under the range $(0,M)$, then there is no incentive to split under perfect information. However, this depends upon the strength of competition. Regardless of how large the split is or how big the rest of the population is, there will always be a strength of competition $\alpha_{r}$ so low that members of a group will choose to split even with perfect information. Solving for that level gives us

$$\alpha_{r}<\frac{2x_{R}}{M}+\frac{1}{M}-\frac{2\sqrt{Mx_{R}+x_{R}^{2}}}{M^{2}}$$

(23)

If this inequality is met, then there will always be group splitting under our assumptions.
